# Asymmetric assembly underlies activation mechanism of neuronal GIRK1/2 channels

**DOI:** 10.64898/2026.10.06.756360

**Authors:** Takeharu Kawano, Liang Wang, Kim W. Chan, Andrew Zorn, Nousheen Parvaiz, Mehreen Zaka, R. Charles Kissell, Carlos Villalba-Galea, Ana Santa-Cruz, Yongcheng Lu, Nihal Aly, James Pentikis, Evangelos Dadiotis, Siyu Li, Meng Cui, Leigh D. Plant, Georgios Skiniotis, Diomedes E. Logothetis

## Abstract

G protein-gated inwardly rectifying potassium (GIRK; Kir3) channels are principal effectors of inhibitory GPCR signaling. Here we combine cryo-electron microscopy, single-molecule imaging, electrophysiology and molecular dynamics to define the architecture and activation of neuronal GIRK1/2 channels. GIRK1/2 preferentially assembles as an asymmetric heterotetramer containing one GIRK1 and three GIRK2 subunits. Three PIP₂ molecules bound to GIRK2 promote partial cytoplasmic-domain docking, whereas engagement of a fourth PIP₂ by GIRK1 completes docking and opens the helix bundle crossing gate. GIRK1 also remodels the electrostatic environment beneath the selectivity filter, where its E141 and D173 residues create a permeation module that enhances K⁺ conduction. The GIRK1-selective activator ML297 binds at the GIRK1–GIRK2 interface and controls both permeation and gating. These findings establish how asymmetric assembly couples sequential lipid-dependent gating to efficient K⁺ permeation and GIRK1-dependent pharmacological activation.

## Main Text

G protein-gated inwardly rectifying potassium (GIRK; Kir3) channels mediate a major inhibitory signaling pathway downstream of Gi/o-coupled G protein-coupled receptors (GPCRs). Activation of receptors including GABA_B_, μ-opioid, dopamine D₂, adenosine A₁ and serotonin 5-HT₁A receptors releases Gβγ subunits that directly activate GIRK channels, producing membrane hyperpolarization and suppression of neuronal excitability. Through this mechanism, GIRK channels regulate synaptic inhibition, cardiac rhythm, hormone secretion and many aspects of nervous system function^1–4^.

Among the four mammalian GIRK subunits, GIRK2 forms functional homotetramers whereas GIRK1 is obligately heteromeric. In the central nervous system, GIRK1/2 heterotetramers constitute the predominant neuronal GIRK channel and exhibit substantially greater basal and agonist-evoked activity than GIRK2 homotetramers. Consequently, GIRK1-containing channels have emerged as attractive therapeutic targets for epilepsy, pain, addiction and other neuropsychiatric disorders.^1, 5–11^ Despite this importance, the molecular mechanisms by which GIRK1 enhances channel function and confers selective pharmacological sensitivity remain poorly understood.

Structural studies of GIRK2 homotetramers have established that phosphatidylinositol-4,5-bisphosphate (PIP₂) is an essential cofactor that promotes docking of the cytoplasmic domain (CTD) onto the transmembrane domain (TMD), leading to opening of the helix bundle crossing (HBC) gate. Intracellular Na⁺, cholesterol and Gβγ further shift the conformational equilibrium toward conducting states^2–4,12–16^. However, virtually all mechanistic insight has come from GIRK2 homotetramers, leaving unresolved how incorporation of GIRK1 modifies channel architecture, activation and pharmacological responsiveness.

Three key mechanistic questions therefore remain. What is the experimentally determined stoichiometry of neuronal GIRK1/2 channels compared to previously proposed models^17,18^? How does GIRK1 alter the structural mechanism of PIP₂-dependent stabilization of the conducting state compared to that in GIRK2 homotetramers^19–21^? And how does GIRK1 confer enhanced channel activity together with sensitivity to urea-based activators such as ML297^22–24^? Resolving these questions is essential for understanding neuronal GIRK1/2 function and for the rational development of subtype-selective modulators.

Here, we combined cryo-electron microscopy, single-molecule total internal reflection fluorescence microscopy, electrophysiology and gating molecular dynamics (GMD) simulations to define the molecular architecture and activation mechanism of neuronal GIRK1/2 channels. We show that GIRK1/2 preferentially assembles as an asymmetric heterotetramer containing one GIRK1 and three GIRK2 subunits. This asymmetric organization establishes sequential PIP₂-dependent activation in which engagement of a fourth PIP₂ molecule by the single GIRK1 subunit completes CTD docking and opens the HBC gate. In addition, incorporation of GIRK1 creates a specialized permeation module beneath the selectivity filter that enhances K⁺ conduction and provides the structural basis for selective pharmacological activation by ML297. Together, these findings identify asymmetric incorporation of a single GIRK1 subunit as the organizing principle that integrates lipid sensing, ion permeation and pharmacological modulation in the principal neuronal GIRK1/2 channel.

## Results

### Neuronal GIRK1/2 channels assemble as asymmetric 1:3 heterotetramers

The molecular architecture of neuronal GIRK1/2 channels has remained uncertain because previous biochemical and functional studies have generally assumed a symmetric heterotetramer containing two GIRK1 and two GIRK2 subunits. To define the stoichiometry and structural organization of the predominant neuronal GIRK channel, we purified recombinant GIRK1/2 complexes (human GIRK1 and mouse GIRK2, which shares 96.2% making it a suitable structural surrogate for the human channel) (Extended Data Fig. 1) and determined four cryo-electron microscopy (cryo-EM) structures representing successive functional states: an undocked ligand-free channel lacking bound PIP₂, a partially docked channel with three PIP₂ molecules bound to the GIRK2 subunits, a fully docked channel with four PIP₂ molecules, and a fully docked ML297-bound complex (Fig. 1a,b; Extended Data Figs. 1–5; Table 1).

**Figure 1.**
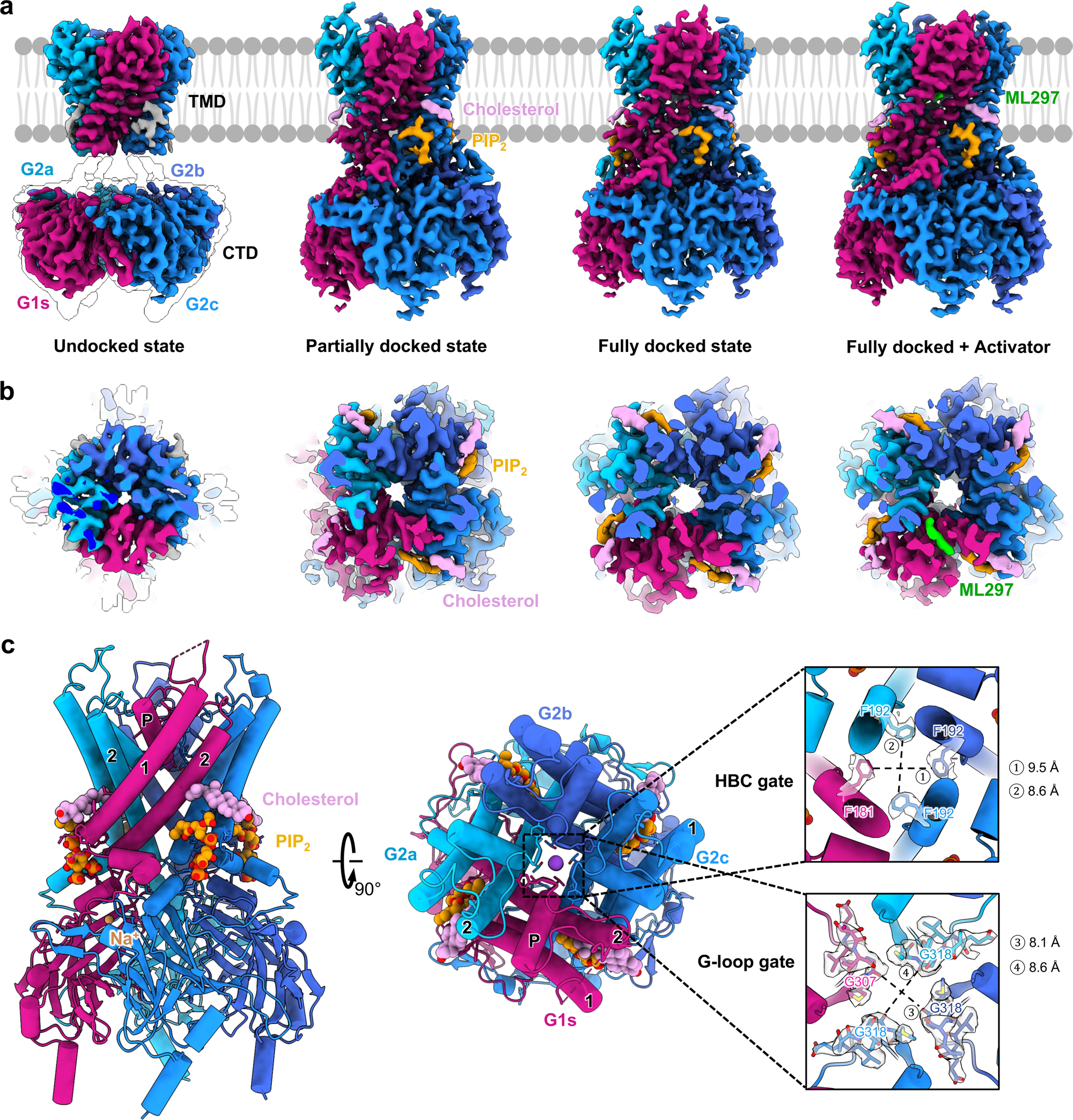
Conformational states and stoichiometry of the GIRK1/2 heterotetramer. a, Cryo-EM reconstructions of GIRK1/2 resolve four distinct conformations that differ in CTD–TMD engagement and ligand occupancy: an undocked state without PIP₂, a partially docked state containing three PIP₂ molecules bound to the GIRK2 subunits, a fully docked state with four PIP₂ molecules, and a fully docked state in the presence of ML297. Each subunit is colored distinctly, with GIRK1 in red and GIRK2 in different shades of blue. b, Top-view sliced cryo-EM density maps of the corresponding states shown in a, viewed from the extracellular side. These sections highlight the asymmetric arrangement of one GIRK1 and three GIRK2 subunits, as well as the state-dependent positions of PIP₂, cholesterol and ML297 within the TMD. c, Model of the fully docked GIRK1/2 heterotetramer bound to four PIP₂, cholesterol and Na⁺, viewed parallel to the membrane (left) and from the extracellular side (middle). The asymmetric arrangement of one GIRK1 and three GIRK2 subunits is evident. Insets show the HBC and G-loop gates and the corresponding minimum heavy-atom distances at each constriction.

**Table 1.**
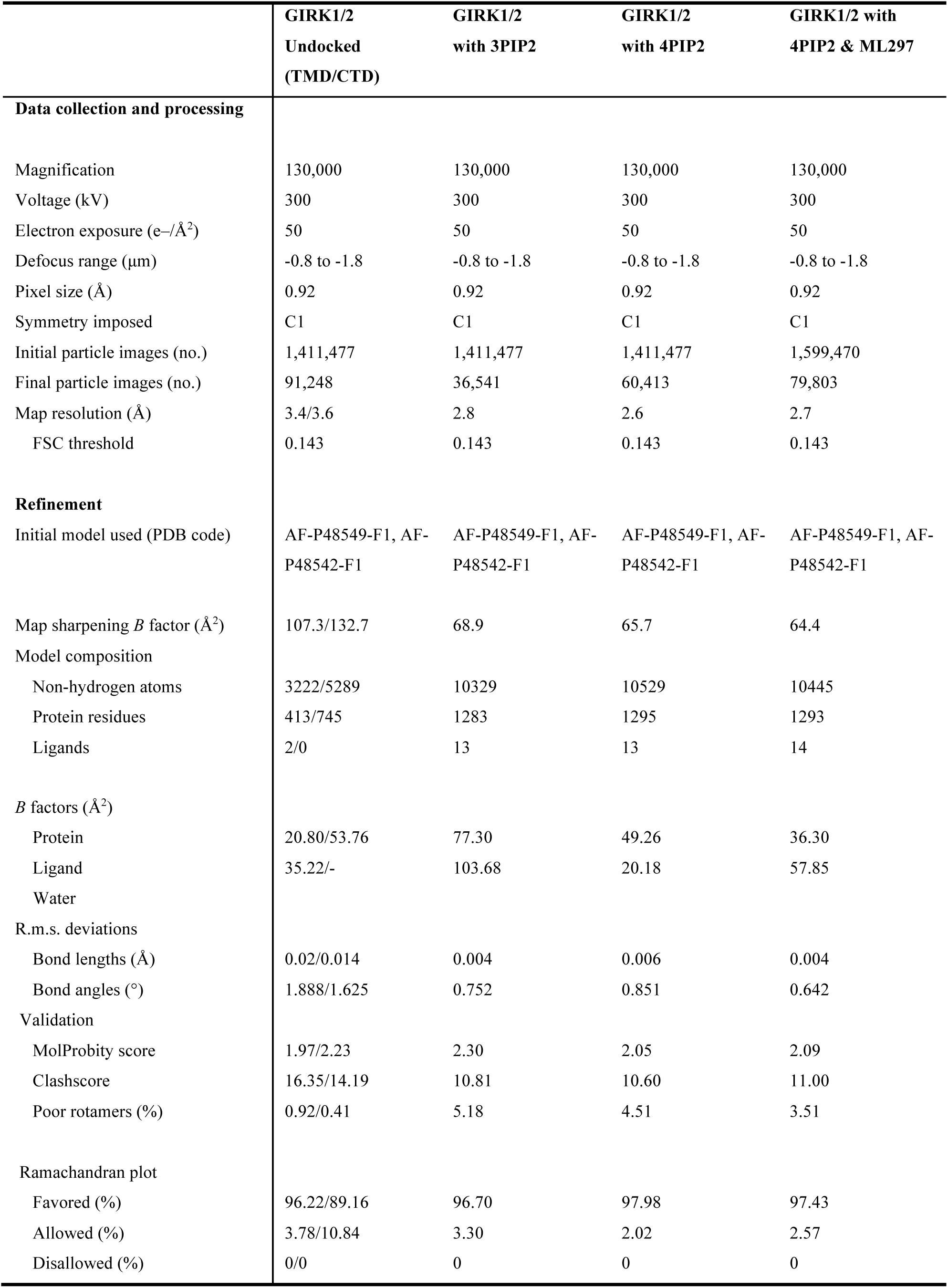
Cryo-EM data collection, refinement and validation statistics.

|  | GIRK1/2<br>Undocked<br>(TMD/CTD) | GIRK1/2<br>with 3PIP2 | GIRK1/2<br>with 4PIP2 | GIRK1/2 with<br>4PIP2 & ML297 |
| --- | --- | --- | --- | --- |
| Data collection and processing |  |  |  |  |
| Magnification | 130,000 | 130,000 | 130,000 | 130,000 |
| Voltage (kV) | 300 | 300 | 300 | 300 |
| Electron exposure (e-/Å<sup>2</sup>) | 50 | 50 | 50 | 50 |
| Defocus range (µm) | -0.8 to -1.8 | -0.8 to -1.8 | -0.8 to -1.8 | -0.8 to -1.8 |
| Pixel size (Å) | 0.92 | 0.92 | 0.92 | 0.92 |
| Symmetry imposed | C1 | C1 | C1 | C1 |
| Initial particle images (no.) | 1,411,477 | 1,411,477 | 1,411,477 | 1,599,470 |
| Final particle images (no.) | 91,248 | 36,541 | 60,413 | 79,803 |
| Map resolution (Å) | 3.4/3.6 | 2.8 | 2.6 | 2.7 |
| FSC threshold | 0.143 | 0.143 | 0.143 | 0.143 |
| Refinement |  |  |  |  |
| Initial model used (PDB code) | AF-P48549-F1, AF-P48542-F1 | AF-P48549-F1, AF-P48542-F1 | AF-P48549-F1, AF-P48542-F1 | AF-P48549-F1, AF-P48542-F1 |
| Map sharpening *B* factor (Å<sup>2</sup>) | 107.3/132.7 | 68.9 | 65.7 | 64.4 |
| Model composition |  |  |  |  |
| Non-hydrogen atoms | 3222/5289 | 10329 | 10529 | 10445 |
| Protein residues | 413/745 | 1283 | 1295 | 1293 |
| Ligands | 2/0 | 13 | 13 | 14 |
| *B* factors (Å<sup>2</sup>) |  |  |  |  |
| Protein | 20.80/53.76 | 77.30 | 49.26 | 36.30 |
| Ligand | 35.22/- | 103.68 | 20.18 | 57.85 |
| Water |  |  |  |  |
| R.m.s. deviations |  |  |  |  |
| Bond lengths (Å) | 0.02/0.014 | 0.004 | 0.006 | 0.004 |
| Bond angles (°) | 1.888/1.625 | 0.752 | 0.851 | 0.642 |
| Validation |  |  |  |  |
| MolProbity score | 1.97/2.23 | 2.30 | 2.05 | 2.09 |
| Clashscore | 16.35/14.19 | 10.81 | 10.60 | 11.00 |
| Poor rotamers (%) | 0.92/0.41 | 5.18 | 4.51 | 3.51 |
| Ramachandran plot |  |  |  |  |
| Favored (%) | 96.22/89.16 | 96.70 | 97.98 | 97.43 |
| Allowed (%) | 3.78/10.84 | 3.30 | 2.02 | 2.57 |
| Disallowed (%) | 0/0 | 0 | 0 | 0 |

All four reconstructions revealed the same asymmetric subunit organization, consisting of a single GIRK1 subunit associated with three GIRK2 subunits (Fig. 1a-c). GIRK1 (denoted as G1s) was readily distinguished from GIRK2 (denoted as G2a, G2b, G2c) by its characteristic extracellular glycosylation, unique sequence features and the presence of the GIRK1-dependent ligand-binding pocket. Importantly, the asymmetric architecture was preserved in every conformational state, indicating that it is an intrinsic property of neuronal GIRK1/2 channels rather than a consequence of channel activation or ligand binding.

The quality of the cryo-EM reconstructions supports an unambiguous assignment of subunit identity. Representative density maps clearly resolved side-chain features that distinguish GIRK1 from GIRK2 (Extended Data Fig. 4). The four reconstructions were independently refined to resolutions ranging from approximately 2.6–3.6 Å, with excellent local resolution throughout both the transmembrane and cytoplasmic domains (Extended Data Fig. 2). Atomic models showed excellent agreement with the experimental density maps (Extended Data Fig. 3), and all reconstructions satisfied standard cryo-EM validation criteria (Table 1). At the HBC gate, G2a and G2c form the narrowest constriction with a minimum distance of 8.6 Å, while at the G-loop gate the narrowest distance of 8.1 Å is formed between G1s-G2b (Fig. 1c).

Because asymmetric stoichiometry was unexpected, we next asked whether it reflected preferential purification or represented the predominant organization of neuronal GIRK1/2 channels in intact cells. We therefore examined channel stoichiometry by single-molecule total internal reflection fluorescence (TIRF) microscopy using GIRK1-TFP and GIRK2-RFP expressed over a broad range of GIRK1:GIRK2 cRNA ratios (Fig. 2a).

**Figure 2.**
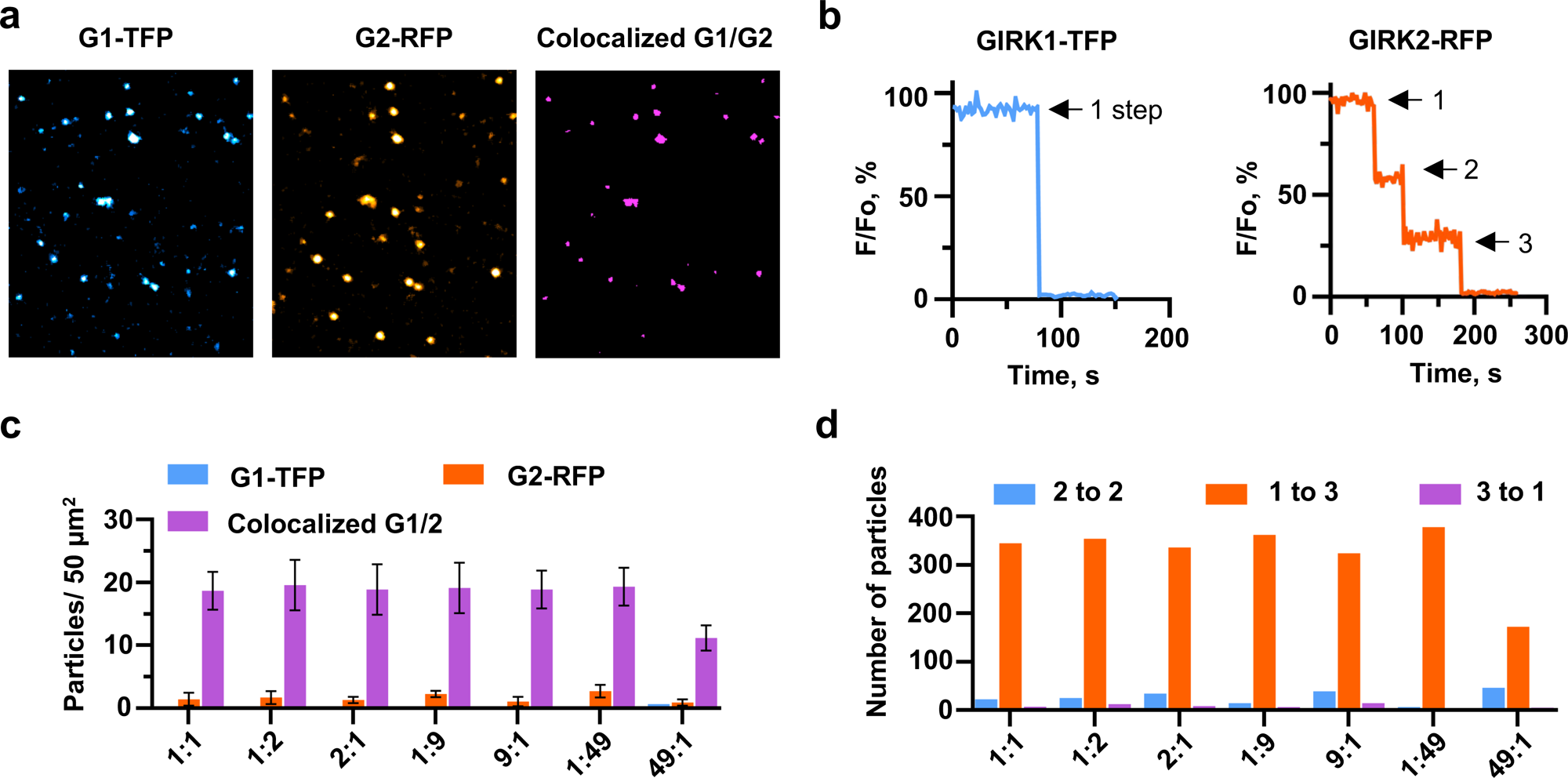
Stoichiometry of the GIRK1/2 tetramer by TIRF microscopy. a, Representative TIRF images from a *Xenopus* oocyte expressing mTFP1-tagged GIRK1 (G1-TFP) and TagRFP-T-tagged GIRK2 (G2-RFP) at a 1:1 cRNA ratio. Colocalized particles are shown in magenta. b, Representative photobleaching records showing one resolved G1-TFP step and three G2-RFP steps in a colocalized particle, consistent with 1:3 stoichiometry. c, Number of G1-TFP, G2-RFP and colocalized particles per 50 μm² at the indicated GIRK1:GIRK2 cRNA injection ratios. Data are from 8–10 oocytes per group from at least three independent experiments. d, Distribution of apparent 2:2, 1:3 and 3:1 subunit compositions among colocalized particles at each injection ratio. The 1:3 population predominated under all conditions examined.

Representative photobleaching traces demonstrated predominantly single-step bleaching for GIRK1-TFP and three-step bleaching for colocalized GIRK2-RFP particles (Fig. 2b), consistent with channels containing one GIRK1 and three GIRK2 subunits. Across expression ratios ranging up to 50-fold in excess of one over the other subunit, colocalized particles remained the predominant membrane population (Fig. 2c), while stoichiometric analysis consistently identified one GIRK1 and three GIRK2 subunits as the major species (Fig. 2d). Statistical analysis of all experiments showed excellent agreement with a 1:3 assembly model (Extended Data Table 1), whereas GIRK2 homotetramers and GIRK1 expressed alone behaved as expected in the corresponding positive and trafficking controls (Extended Data Fig. 6; Extended Data Table 1).

The remarkable agreement between cryo-EM and single-molecule photobleaching demonstrates that asymmetric 1:3 assembly is not an artifact of purification, reconstruction or expression conditions but instead represents the preferred organization of neuronal GIRK1/2 channels. The observation that a single GIRK1 subunit occupies a unique position within an otherwise GIRK2-like tetramer immediately suggested that GIRK1 contributes a specialized structural role rather than simply replacing one of four equivalent subunits.

Compared to the GIRK2 homotetramer^15^ the GIRK1/2 heterotetramer deviates from fourfold symmetry in the transmembrane domain (TMD) relative to the cytoplasmic domain (CTD) by a ∼2 Å displacement (Extended Data Fig. 7). Superposition of the GIRK1 and GIRK2 subunits revealed prominent structural differences in the TMD with the TM2 helices bending in opposite directions as well as in the turret and C-terminal helical regions, in contrast to the highly conserved architecture of the GIRK2 subunits (Extended Data Fig. 8). These observations suggested that asymmetric incorporation of GIRK1 may establish a specialized allosteric module with functional consequences extending beyond channel assembly, prompting us to examine how the unique GIRK1 subunit modifies channel activity.

### A single GIRK1 subunit remodels the permeation pathway of the heterotetramer

Having established that neuronal GIRK1/2 channels preferentially assemble as asymmetric 1:3 heterotetramers, we next asked whether incorporation of a single GIRK1 subunit produces structural specializations that could account for the enhanced activity of GIRK1-containing channels. Comparison of the fully docked GIRK1/2 structure with previously determined GIRK2 homotetramers revealed striking differences immediately beneath the selectivity filter despite the otherwise highly conserved architecture of the pore.

Electrostatic surface calculations showed that incorporation of GIRK1 substantially remodels the electrostatic landscape beneath the selectivity filter (Fig. 3a,b). Whereas the three GIRK2 subunits retain the electrostatic organization previously described for GIRK2 homotetramers, the G1s subunit creates a markedly more electronegative environment extending into the inner permeation pathway. Importantly, this redistribution occurs without detectable changes in the selectivity filter itself, suggesting that GIRK1 modifies the energetic environment through which K⁺ ions pass rather than the ion-coordination sites responsible for selectivity.

**Figure 3.**
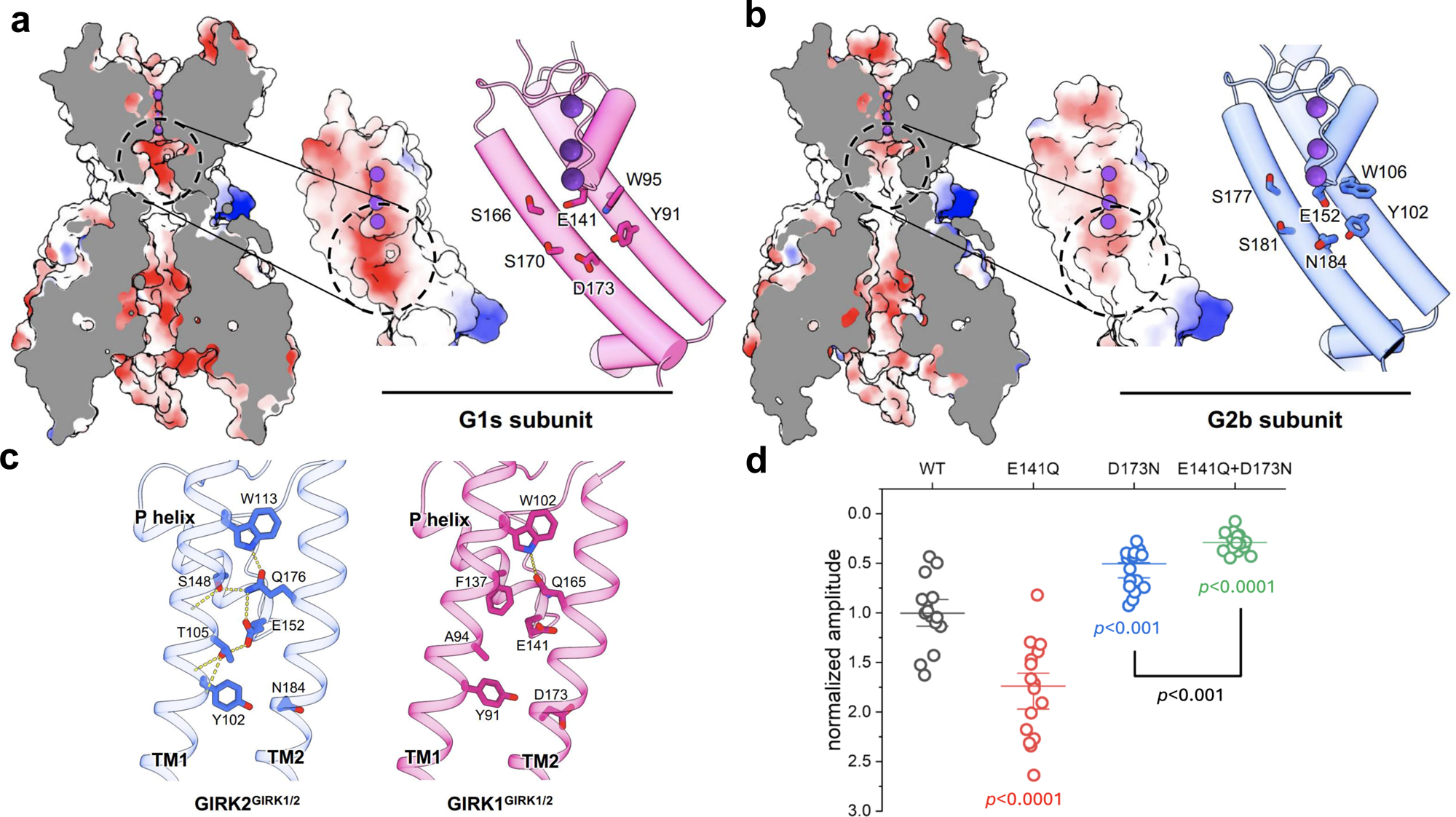
GIRK1 generates a more negative inner-pore electrostatic environment. a,b, Electrostatic surface potentials of GIRK1 (G1s) and GIRK2 (G2b) within the GIRK1/2 heterotetramer (color scale, −10 kT/e in red to +10 kT/e in blue). Enlarged views highlight acidic and polar residues beneath the selectivity filter. c, Local interactions in the pore-helix/TM1/TM2 region of GIRK2 and GIRK1. Yellow dashed lines indicate hydrogen bonds. In GIRK2, interactions among S148.2, Q176.2, T105.2 and E152.2 stabilize an outward orientation of E152.2. The corresponding sites in GIRK1 (F137.1, Q165.1, A94.1 and E141.1) lack this network, allowing E141.1 to face the pore lumen. d, Basal currents from oocytes expressing G1 + G2 or the indicated G1 E141.1/D173.1 mutants with G2. P values relative to G1 + G2 are indicated.

The structural basis for this electrostatic remodeling is revealed by comparison of the hydrogen-bonding networks that position acidic residues beneath the selectivity filter (Fig. 3C). In GIRK2, E152.2 is restrained by an extensive hydrogen-bond network involving S148.2, T105.2 and Q176.2 together with neighboring aromatic residues Y102.2 and W113.2, orienting its side chain away from the permeation pathway. In GIRK1, the corresponding hydrogen-bond donors are replaced by hydrophobic residues, disrupting this network and permitting E141.1 to rotate toward the pore lumen. The adjacent residue D173.1 occupies a position immediately below E141.1, placing two acidic side chains in close proximity and creating a distinct electrostatic environment beneath the selectivity filter that is unique to the asymmetric GIRK1/2 channel.

This structural analysis suggested that E141.1 and D173.1 contribute to channel function by modifying ion permeation rather than by directly altering the principal PIP₂-dependent gating transitions. To test this hypothesis, we examined the effects of mutations predicted to disrupt the GIRK1-specific permeation module. Neutralization of E141.1 (E141.1Q) increased basal current, whereas D173.1N markedly reduced activity, and the double mutant produced a further reduction in current (Fig. 3d; Extended Data Fig. 9). Importantly, total internal reflection fluorescence microscopy demonstrated that these and related mutants exhibited surface expression comparable to wild-type channels (Extended Data Fig. 10), indicating that the altered currents primarily reflect changes in channel function rather than membrane trafficking.

To determine whether disruption of the GIRK2 hydrogen-bond network accounts for the unique properties of GIRK1, we restored individual GIRK2-like interactions within GIRK1. Although mutagenesis of the unique GIRK1 residues that disrupt h-bonding, F137.1S and A94.1T individually, did not reduce activity, their combination, as well as incorporation of D173.1N, significantly impaired channel activity (Extended Data Fig. 9). Conversely, consistent with the established importance of the corresponding pore-helix position in GIRK2, substitution of S148.2 with phenylalanine increased basal GIRK2 activity (Extended Data Fig. 9). Together, these mutational analyses support the structural prediction that disruption of the GIRK2 hydrogen-bond network contributes to the enhanced functional properties conferred by GIRK1.

We next used gating molecular dynamics (GMD) simulations —molecular dynamics simulations designed to relate conformational changes to ion permeation events²⁹— to examine how these experimentally established structural differences could influence ion permeation. Simulations comparing GIRK1/2, GIRK2 and GIRK2(S148F,N184D) predicted that disruption of the GIRK2 hydrogen-bond network together with introduction of an acidic residue at the D173.1-equivalent position favors K⁺ permeation through the inner pore (Extended Data Fig. 11). Thus, the simulations provide a mechanistic framework that is consistent with both the cryo-EM structures and the functional effects of the corresponding mutations.

Together, these structural, functional and computational analyses identify a previously unrecognized permeation module created by asymmetric incorporation of the single GIRK1 subunit. Rather than directly modifying the canonical gating machinery, this module remodels the electrostatic environment immediately beneath the selectivity filter, thereby providing a mechanistic explanation for the enhanced permeation properties of neuronal GIRK1/2 channels.

### Sequential PIP₂ binding drives complete activation of the asymmetric GIRK1/2 channel

The asymmetric architecture of GIRK1/2 suggested that the single GIRK1 subunit might contribute uniquely to channel activation. To define the structural sequence underlying activation, we compared the three ligand-free cryo-EM structures representing successive stages of PIP₂ occupancy: an undocked channel lacking bound PIP₂, a partially docked channel with three PIP₂ molecules bound to the GIRK2 subunits, and a fully docked channel containing a fourth PIP₂ molecule bound to GIRK1 (Figs. 1a,b). These structures capture a progressive activation pathway in which sequential lipid binding drives distinct conformational transitions.

Superposition of the three conformational states revealed progressive outward displacement of the inner TM2 helices accompanied by an approximately 2 Å upward translation of the CTD toward the membrane (Figs. 1b, 4a). The largest conformational rearrangements occur within the GIRK1 subunit, whereas the three GIRK2 subunits undergo comparatively modest changes. The fully occupied state, in which a fourth PIP₂ molecule is bound to GIRK1, is associated with complete CTD docking and widening of the helix bundle crossing (HBC) gate. In contrast, the G-loop gate narrows during the transition from the undocked to the partially docked state and changes only modestly following engagement of the fourth PIP₂ molecule (Fig. 4a). These observations indicate that the final lipid-binding event primarily stabilizes complete CTD docking and HBC opening rather than producing large conformational changes at the G-loop gate.

**Figure 4.**
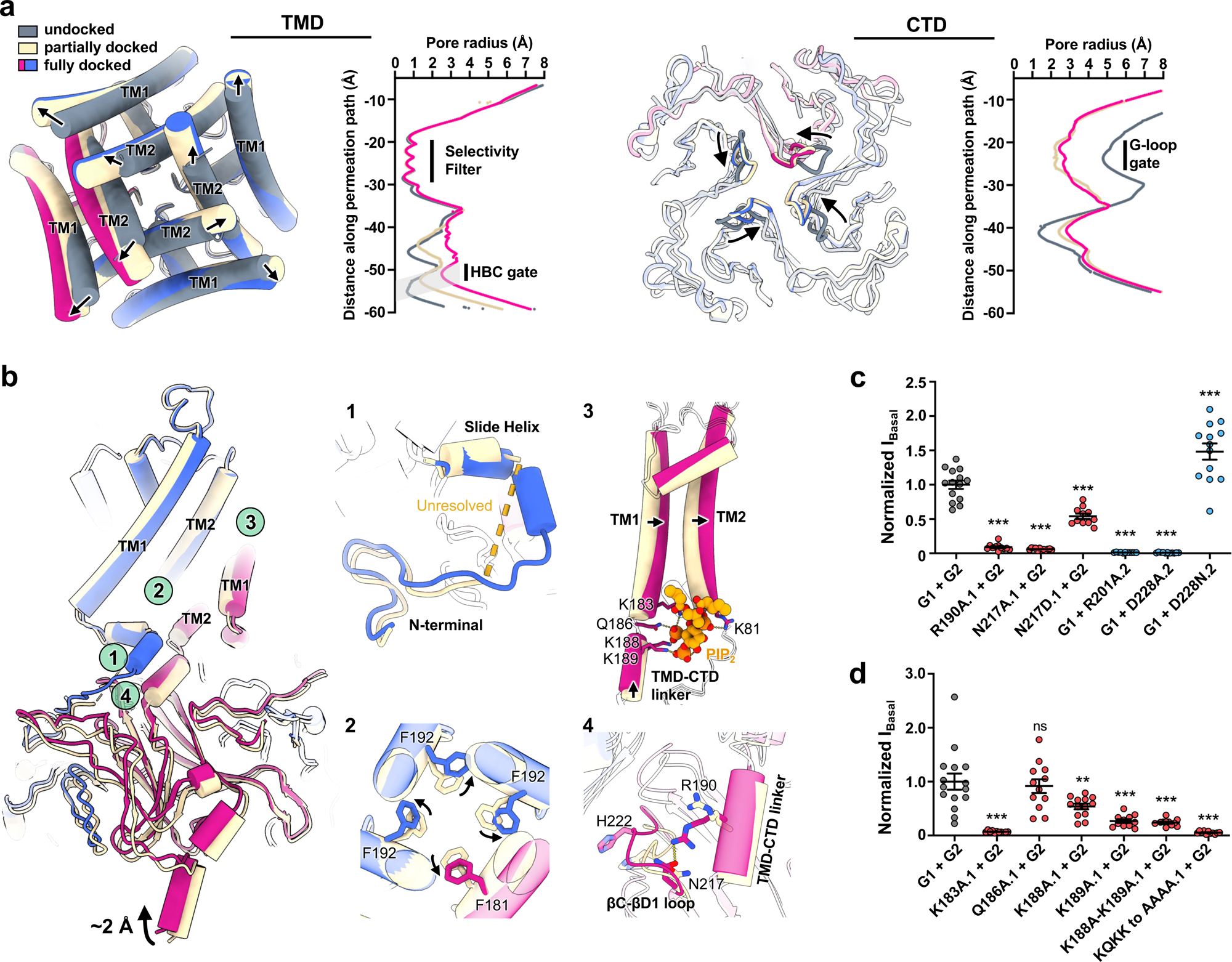
Structural and functional consequences of partial versus full PIP₂ occupancy in GIRK1/2. a, Undocked, three-PIP₂-bound and four-PIP₂-bound GIRK1/2 structures viewed from the cytoplasmic side for the TMD and from the TMD toward the CTD. Superposed TMDs and pore-radius profiles show progressive HBC widening with increasing PIP₂ occupancy; CTD pore-radius profiles show the corresponding G-loop changes. b, Effects of alanine substitution of GIRK1 PIP₂-coordinating residues on basal G1 + G2 currents measured by TEVC in Xenopus oocytes expressing the indicated constructs (0.5 ng cRNA per subunit), 2 d after injection. P values are relative to G1 + G2: **P < 0.01; ***P < 0.001. c, Comparison of the three- and four-PIP₂-bound states, showing ∼2 Å upward CTD displacement upon full PIP₂ occupancy. Insets show (1) unresolved slide-helix density in the three-PIP₂-bound state, (2) differences at the HBC constriction, (3) PIP₂-coordinating residues in GIRK1 and (4) the R190.1–N217.1 interaction observed in the four-PIP₂-bound state. d, Effects of mutations at interactions associated with binding of the fourth PIP₂ molecule on basal G1 + G2 currents. Experiments were performed as in B. ***P < 0.001 relative to G1 + G2; ns, not significant.

Completion of PIP₂ occupancy also orders structural elements that remain flexible in the partially docked state. Prior to full lipid occupancy, the GIRK1 slide helix exhibits increased conformational mobility (Fig. 4c, panel 1). Binding of the fourth PIP₂ stabilizes interactions linking the slide helix, the TMD–CTD linker and the membrane interface (Fig. 4c, panels 2–4), while simultaneously promoting outward rotation of the phenylalanine side chains that form the HBC gate, increasing the minimum pore diameter to dimensions compatible with K⁺ permeation (Figs. 4a,c).

To determine whether the interactions identified in the cryo-EM structures contribute to channel activation, we examined residues coordinating PIP₂ within the GIRK1 lipid-binding pocket (Fig. 4b). Neutralization of individual coordinating residues progressively reduced basal activity, with K183.1 producing the largest effect, followed by K189.1 and K188.1, whereas simultaneous disruption of all four coordinating residues nearly abolished channel function. These functional effects closely parallel the structural importance of the GIRK1 PIP₂-binding site in stabilizing the fully docked conformation.

We next tested interactions that become established only after engagement of the fourth PIP₂ molecule by G1s. Mutations disrupting these contacts reduced basal channel activity, whereas D228N.2, predicted to destabilize an inhibitory interaction^28^, enhanced basal current (Fig. 4d). Likewise, the R190.1–N217.1 interaction identified in the fully docked structure (Fig. 4c, panel 4) proved important for basal channel activity (Fig. 4d). Thus, residues that stabilize the fully docked conformation identified structurally are also required for efficient channel activation.

Together, these structural and functional analyses support a sequential activation mechanism in which binding of three PIP₂ molecules to the GIRK2 subunits initiates CTD docking, whereas engagement of the fourth PIP₂ molecule by the single G1s subunit completes CTD docking and stabilizes opening of the HBC gate. This mechanism explains why asymmetric incorporation of GIRK1 introduces a unique final activation step that cannot occur in GIRK2 homotetramers.

Importantly, these observations distinguish regulation of channel gating from regulation of ion permeation. The principal PIP₂-dependent conformational changes localize to the TMD–CTD interface and HBC gate (Figs. 1, 4), whereas mutations within the GIRK1 permeation module predominantly alter macroscopic current without directly affecting these gating transitions. Together, the structural, functional and computational data therefore support mechanistically distinct, but tightly coupled, contributions of gating and permeation to neuronal GIRK1/2 channel activity.

### ML297 selectively engages the GIRK1 permeation module to potentiate channel activation

The structural and functional analyses above indicate that asymmetric incorporation of GIRK1 confers two mechanistically distinct properties upon the heterotetramer: formation of a specialized permeation module beneath the selectivity filter and completion of the final PIP₂-dependent activation step. We therefore asked how the GIRK1-selective activator ML297 exploits these unique structural features to potentiate channel activity.

To define the structural basis of selective activation, we determined the cryo-EM structure of GIRK1/2 bound to ML297 at 2.7 Å resolution. ML297 occupies a well-defined allosteric pocket formed between TM1 of the GIRK1 subunit and TM2 of the adjacent GIRK2 subunit (Fig. 5a). The high-quality density permitted unambiguous modeling of the urea scaffold together with its pyrazole and aromatic substituents, revealing an extensive network of hydrogen-bonding and hydrophobic interactions that explains the preference of ML297 for GIRK1-containing heterotetramers.

**Figure 5.**
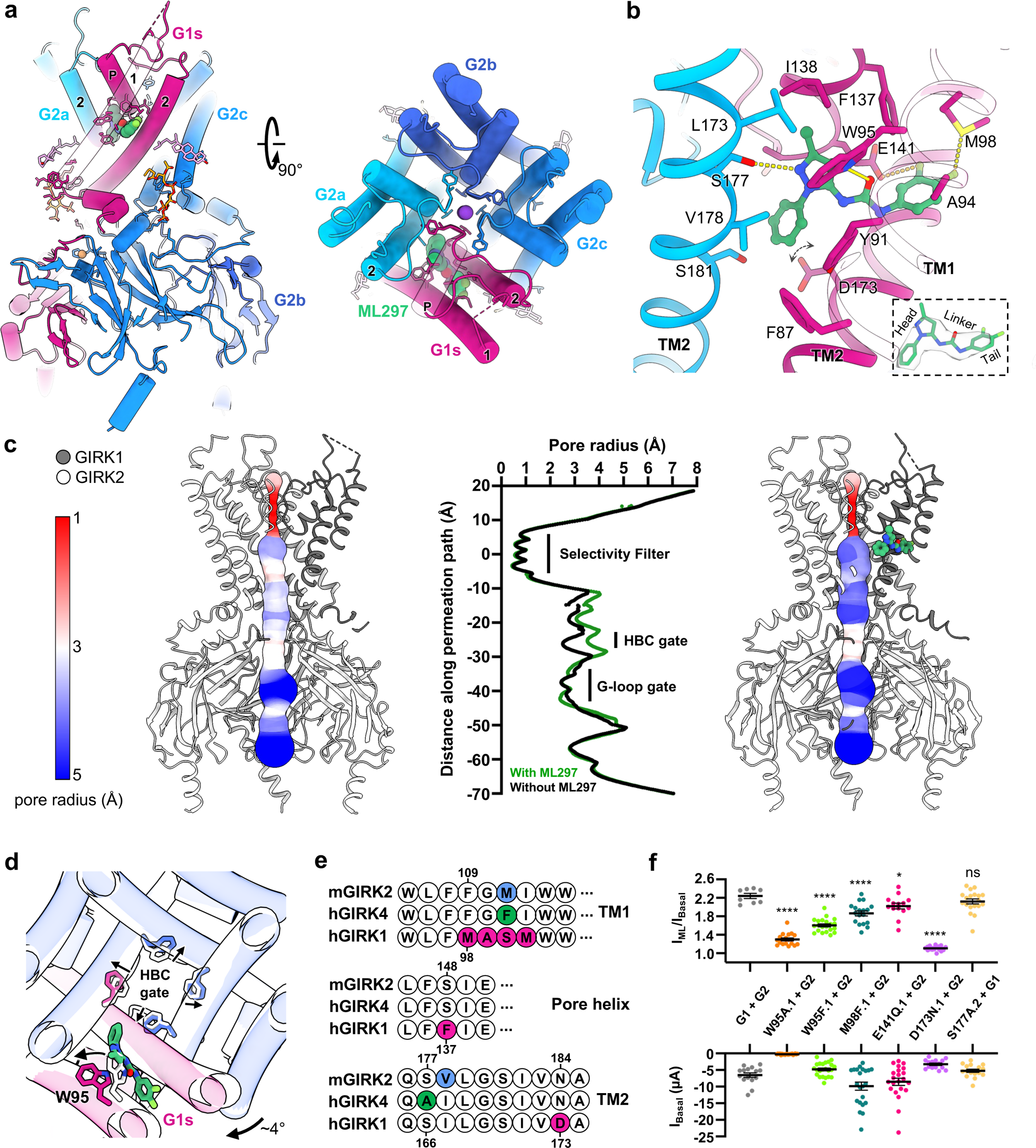
ML297 binds an asymmetric transmembrane pocket and promotes HBC expansion. a, Cryo-EM structure of ML297-bound GIRK1/2 viewed parallel to the membrane (left) and from the extracellular side (right). ML297 occupies a pocket formed by GIRK1 (G1s) and the neighboring GIRK2 subunit. b, ML297-binding site. Interacting residues are highlighted and hydrogen bonds are shown as yellow dashed lines. The inset shows cryo-EM density for ML297 and the head, linker and tail regions. Double-headed arrows indicate conformational mobility of the D173.1 side chain. c, Pore-radius profiles and pore-geometry representations of GIRK1/2 with and without ML297. ML297 widens the pathway at the HBC, whereas the selectivity filter and G-loop gate remain. d, Structural comparison of GIRK1/2 with and without ML297. ML297 binding is associated with outward displacement of GIRK1 W95.1 and rearrangement of surrounding TM1/TM2 helices, producing local expansion near the HBC and a small clockwise TMD rotation. e, Sequence alignment of mouse GIRK2, human GIRK4 and human GIRK1 highlighting residues at positions forming the ML297-binding pocket. f, ML297 activation, expressed as current in 10 μM ML297 divided by basal current (I_ML_/I_basal_), and basal current in *Xenopus* oocytes expressing G1 + G2 with the indicated GIRK1 or GIRK2 mutations. Data are from n = 9–21 oocytes per group across N = 3 independent experiments. Statistical comparisons are with wild-type G1 + G2 using two-tailed unpaired Welch’s t-tests; *P < 0.05, ****P < 0.0001; ns, not significant.

Comparison of the fully docked ligand-free and ML297-bound structures revealed that ML297 does not induce a fundamentally new conformation. Rather, ligand binding further stabilizes the PIP₂-bound conducting state by promoting additional outward displacement of the TM2 helices and modest widening of the helix bundle crossing (HBC), while preserving the overall architecture of both the selectivity filter and cytoplasmic domain (Fig. 5b,c). The pore-radius analysis further illustrates that ML297 primarily expands the intracellular HBC constriction while leaving the selectivity filter essentially unchanged (Fig. 5c), indicating that the ligand potentiates an existing activation pathway rather than introducing an alternative gating mechanism.

Closer inspection of the binding pocket provides a structural explanation for GIRK1 selectivity (Fig. 5b,d). The urea core bridges residues contributed by GIRK1 and the neighboring GIRK2 subunit, creating an interaction network that can arise only within the asymmetric heterotetramer. Particularly noteworthy is the unresolved dynamic interaction with GIRK1 residue D173.1, which lies within the permeation module identified above, together with neighboring residues lining the adjacent GIRK2 subunit (Fig. 5d). The pyrazole substituent also occupies positions compatible with interactions with S177.2, one helical turn above the G180.2 hinge, suggesting that ligand binding samples multiple conformations within the asymmetric interface while maintaining the same overall binding mode.

To determine whether these structural interactions contribute to channel activation, we mutated residues predicted to contact ML297 (Fig. 5e,f). Disruption of multiple pocket residues altered agonist-evoked currents, supporting the structural assignments. Among these, neutralization of D173.1 produced the most pronounced reduction in ligand efficacy despite normal surface expression (Extended Data Fig. 10), identifying this residue as a critical determinant of pharmacological activation. Whereas several mutations altered basal activity while retaining partial responses to ML297, D173.1N almost completely abolished agonist potentiation (Fig. 5f). Because D173.1 lies outside the principal PIP₂-binding network responsible for CTD docking and HBC opening, the marked loss of ligand efficacy is unlikely to result solely from impaired PIP₂-dependent gating.

Given the importance of D173.1 in ML297-induced activity and the fact that it could not be resolved in the structure (Fig. 5b) due to its flexibility, we used GMD simulations to investigate how these experimentally identified structural elements may be dynamically coupled during channel activation (Extended Data Fig. 12). Throughout the simulations, the ligand dynamically engaged residues within GIRK1 TM1 (W95.1), the pore helix (E141.1), GIRK1 TM2 (D173.1) and GIRK2 TM2 (S177.2 or S181.2 on opposite sides of the G180.2 hinge), stabilizing an interaction network that spans the asymmetric GIRK1–GIRK2 interface (Extended Data Fig. 12). This local binding stabilization propagated through the transmembrane domain was associated with persistence of a more widely open HBC configuration (Fig. 6), consistent with the structural differences observed between the ligand-free and ML297-bound cryo-EM structures (Fig. 5b,c). Yet, the GMD simulations resulted in modest increases in ion permeation over unliganded channels, perhaps due to the short 0.5 µs simulation runs.

**Figure 6.**
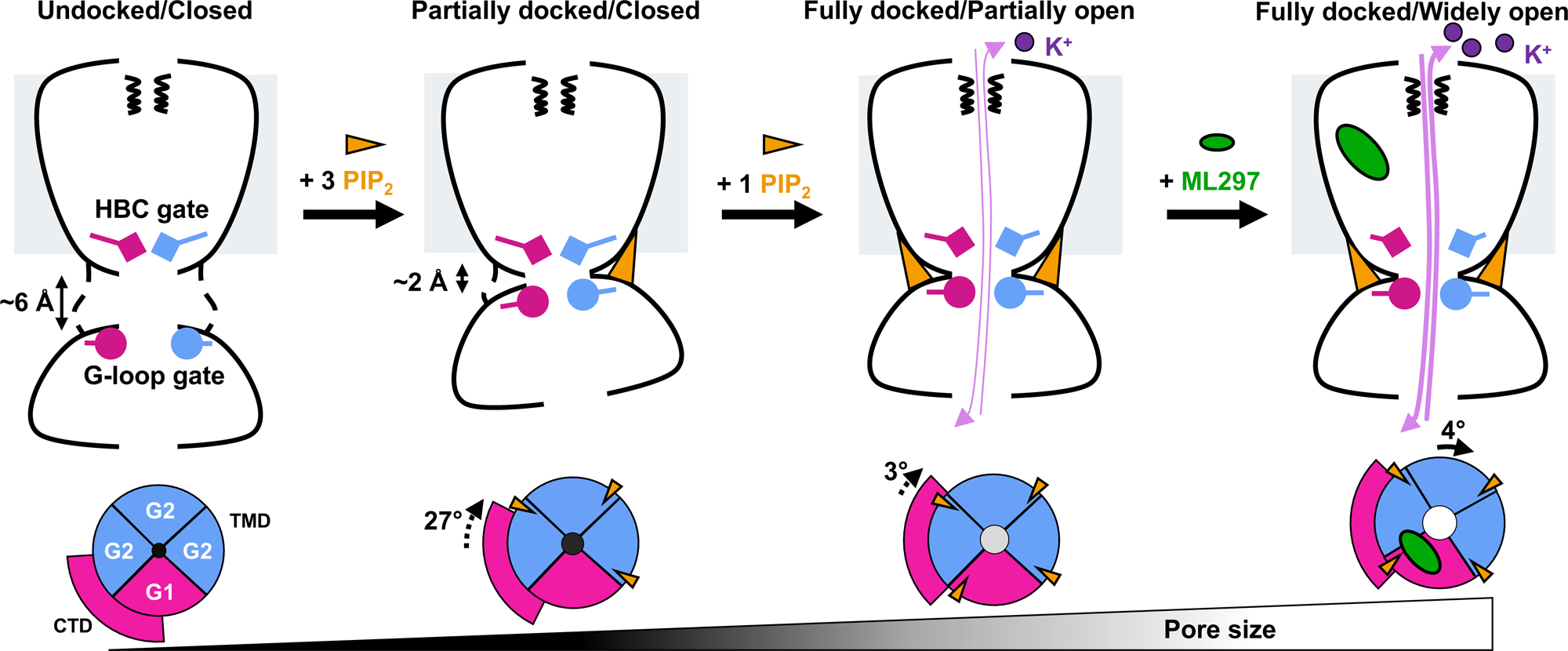
Sequential conformational transitions of GIRK1/2 driven by PIP₂ and ML297. In the undocked state, no PIP₂ is resolved, the CTD is displaced from the TMD by ∼6 Å and the HBC is closed, whereas the G-loop gate is widest. Binding of three PIP₂ molecules to the GIRK2 subunits produces partial CTD docking and ∼27° clockwise CTD rotation, while the GIRK1 side remains ∼2 Å disengaged and the HBC remains closed. Binding of the fourth PIP₂ molecule at GIRK1 completes CTD docking, adds ∼3° clockwise rotation and partially opens the HBC to support basal K⁺ permeation. ML297 binding is associated with an additional ∼4° clockwise TMD twist and further HBC widening, increasing K⁺ conduction. Dashed and solid arrows indicate CTD and TMD rotations, respectively.

Together, the structural, functional and computational analyses support a model in which ML297 exploits the specialized architecture created by asymmetric incorporation of a single GIRK1 subunit. Rather than inducing a novel open state, ML297 stabilizes the fully PIP₂-bound conducting conformation while simultaneously engaging the GIRK1 permeation module through D173.1 and neighboring residues. Thus, ML297-induced pharmacological activation emerges from coordinated effects on both permeation and gating, rather than modulation of either process in isolation.

## Discussion

Cryo-EM and single-molecule photobleaching independently demonstrated that neuronal GIRK1/2 channels preferentially assemble as asymmetric complexes containing one GIRK1 and three GIRK2 subunits. This architecture establishes the organizing principle underlying the unique functional properties of neuronal GIRK channels. The single GIRK1 subunit contributes three distinct mechanistic features to the heterotetramer. It completes the final PIP₂-dependent activation step, remodels the permeation pathway beneath the selectivity filter, and creates an allosteric binding pocket exploited by GIRK1-dependent activators. Thus, asymmetric incorporation of GIRK1 creates a specialized allosteric module that integrates lipid sensing, ion permeation and pharmacological modulation.

More broadly, these findings suggest that heteromeric ion channels through incorporation of a single specialized subunit can generate new allosteric functions unavailable to homomeric channels, providing an efficient evolutionary strategy for expanding functional diversity without increasing oligomeric complexity. Similar principles may underlie functional specialization in other heteromeric channel families³¹˒³².

Our structures further provide a structural framework for understanding how PIP₂ progressively activates neuronal GIRK1/2 channels. Rather than acting as a binary on-off switch, PIP₂ binding proceeds sequentially. Figure 6 summarizes in a cartoon model the sequential conformational transitions of GIRK1/2 driven by PIP_2_. Occupancy of the three GIRK2 binding sites initiates a large clockwise rotation of the CTD relative to TMD and its partial docking to the TMD. Engagement of the fourth PIP_2_ on GIRK1 completes an additional small clockwise rotation of the CTD relative to the TMD and its full docking stabilizing opening of the HBC gate. The resulting series of conformational states suggests that asymmetric incorporation of GIRK1 introduces a final activation step that is absent from GIRK2 homotetramers. Equally importantly, the GIRK1-specific hydrogen-bond network remodels the electrostatic environment beneath the selectivity filter without substantially altering the canonical gating machinery. Together, the structural, mutagenesis and GMD analyses support the view that efficient K⁺ conduction depends not only on opening of the intracellular gate but also on optimization of the energetic landscape through which ions traverse the inner pore.

The structural basis for GIRK1-dependent activation has remained incompletely understood despite extensive medicinal chemistry centered on ML297 and related urea derivatives. Our findings indicate that subtype selectivity arises not simply because ML297 binds GIRK1, but because it recognizes an asymmetric interface that exists only after heteromer assembly. Thus, the pharmacological binding pocket is itself a consequence of asymmetric architecture. Ligand efficacy reflects coordinated stabilization of both gating and permeation. ML297 enhances HBC opening within the fully PIP₂-bound channel while simultaneously engaging residues within the GIRK1 permeation module, particularly D173.1. The pronounced loss of agonist efficacy following D173.1 neutralization, despite preservation of the principal PIP₂-dependent gating machinery, indicates that optimization of ion permeation contributes substantially to pharmacological activation. ML297 is therefore better viewed as stabilizing a highly conducting state than as simply increasing the probability of channel opening. Differences in the neighboring GIRK subunit may further influence subtype selectivity. The replacement of GIRK2 S177.2 by GIRK4 A172.4, together with differences in interactions with residues surrounding the conserved glycine hinge, may alter coupling between ligand binding, TM2 flexibility and HBC opening. This hypothesis provides a structural framework for understanding the divergent effects of ML297 on GIRK1/2 and GIRK1/4 channels and suggests testable strategies for designing next-generation subtype-selective modulators³³.

One of the broader implications of this work is that structural asymmetry can create new opportunities for selective pharmacological intervention. Drug discovery for heteromeric ion channels has traditionally focused on highly conserved orthosteric or pore-forming regions that offer limited opportunities for subtype discrimination. In contrast, asymmetric assembly generates interfaces that exist only within specific heteromeric combinations and therefore provide unique structural environments for selective ligand recognition. The GIRK1/2 channel illustrates this principle particularly clearly. Incorporation of a single GIRK1 subunit simultaneously creates a unique activation mechanism, a specialized permeation module and an asymmetric allosteric pocket that can be preferentially targeted by small molecules. Similar strategies may prove broadly applicable to other heteromeric ion channels that have thus far resisted subtype-selective pharmacological modulation.

## Online Methods

### Molecular biology and biochemistry

GIRK1/2 (mGIRK2-TEV-FLAG-p2A-hGIRK1-TEV-GFP-His10) was constructed in the pFastBac vector for expression in Sf9 insect cells. For mammalian expression, GIRK1/2 (mGIRK2-TEV-FLAG-p2A-hGIRK1-TEV-GFP-His10) was subcloned into pEG_BacMam vector^34^ by GenScript (Piscataway, NJ). The baculoviruses were generated using the Bac-to-Bac system (Thermo Fisher Scientific, Waltham, MA).

### Electrophysiology

The pMAX(+) (in-house vector) and pXoom^35^ vectors were used for expression in HEK293T cells and Xenopus oocytes. The pGEMHE^36^ vector was also used for Xenopus oocytes. Wild-type and mutant constructs of GIRK1 and GIRK2 were in pXoom or pGEMHE. Mutations were introduced by QuikChange site-directed mutagenesis (Agilent, Santa Clara, CA) and confirmed by sequencing. For TEVC using Xenopus oocytes, mRNAs were transcribed using mMESSAGE mMACHINE T7 Transcription Kit (Thermo Fisher Scientific, Waltham, MA) following plasmid linearization. TIRF microscopy: GIRK1-HA-mTFP1 and GIRK2-FLAG-TagRFP-T in pMAX(+) were ordered from GenScript (Piscataway, NJ).

### Protein Expression

GIRK1/2 channels were expressed in HEK293S GnTI - cells or insect Sf9 cells using baculoviruses generated in-house. Suspension cultures of HEK293S GnTI-cells were maintained at a density of 0.2’ 10^6^ cells/mL at 140 rpm and 37 °C with 8% CO2. When cultures reached a density of 2-3.5’ 10^6^ cells/mL, P4 baculovirus encoding the channel of interest was added and incubated for overnight. On the next day, sodium butyrate was added to a final concentration of 10 mM, and the incubation temperature was reduced to 30°C while maintaining 8% CO2. Cultures were incubated for an additional 48 h. The cells were then harvested by centrifugation at 8,000 g for 10 min at 4°C. Sf9 cells were cultured in suspension with shaking at 140 rpm in a 27 °C orbital shaker incubator. GIRK1/2 baculoviruses were added to Sf9 cells at a density of 2-2.5’ 106 cells/mL. Cells were harvested 48 h later by centrifugation. <u>Membrane Preparation:</u> Induced cell pellets were resuspended in ice-cold lysis buffer containing 50 mM Tris-HCL (pH 8), 200 mM NaCl, 20 mM KCl, 1 mM EDTA, 10% (v/v) glycerol and 1 mM PMSF, supplemented with a fresh protease inhibitor cocktail (Thermo Scientific, Cat. No. A32963). Cells were lysed with a microfluidizer at 4000 psi at 4°C. Cell lysate was centrifuged at 35,000 rpm for 30 min in a 45 Ti rotor (Beckman Coulter) to remove soluble protein fractions. The resultant pellet was resuspended in wash buffer containing 50 mM Tris-HCL (pH8), 500 mM KCl, 1 mM EDTA, 10% glycerol, 1 mM PMSF, and centrifuged again at 35,000 rpm for 30 min in a 45 Ti rotor (Beckman Coulter). The final membrane pellets were resuspended in solubilization buffer containing 50 mM Tris-HCL (pH 8), 150 mM KCl, 0.1 mM PMSF, supplemented with fresh protease inhibitors. Membrane protein concentration was determined using a BCA assay (Thermo Scientific, Cat. No. A55864). Membranes were either flash-frozen in liquid nitrogen and stored at -80°C or immediately used for detergent extraction. <u>Detergent Extraction of Mammalian Cell Membranes and Purification:</u> Membranes were diluted to 5 mg/mL in solubilization buffer and extracted with 1% (w/v) n-dodecyl-β-D-maltoside (DDM, Gold Biotechnology, Cat. No. DDM10) for 2 h at 4°C. The mixture was centrifuged at 18,600 rpm for 20 min in a 45 Ti rotor (Beckman Coulter). The supernatant containing solubilized membrane proteins was collected and incubated with Ni-NTA agarose beads (GoldBio, Cat. No. H-320) for 2 h to overnight at 4°C. The resin was washed with 10 column volumes (CV) of wash buffer containing 50 mM Tris-HCl (pH 8), 150 mM KCl, 10 mM imidazole, 0.1% DDM, 20% glycerol, 0.1 mM PMSF. Bound proteins were eluted using the wash buffer containing 400 mM imidazole. The imidazole elution was incubated Anti-DYKDDDDK G1 Affinity Resin (GenScript, Cat. No. L00432) for 2 h at 4°C. The resin was subjected to 10 column volumes (CV) of wash buffer containing 20 mM Tris-HCl (pH 8), 150 mM KCl, 200 mM NaCl, 0.05% GDN, 20% glycerol, 0.1 mM PMSF, and 0.5 mM EDTA. GIRK1/2 proteins were eluted using the same buffer supplemented with 200 µg/mL DYKDDDDK-peptide (APExBIO, Cat. No. A6002). GIRK1/2 proteins were TEV-cleaved at 4 °C overnight and purified by Size Exclusion Chromatography (SEC) using Superose6 Increase 10/300 GL (Cytiva, Cat. No. 29091596) column pre-equilibrated with 20 mM Tris-HCl (pH 7.4), 100 mM KCl, 40 mM NaCl, 0.005% GDN, 1 mM DTT, and 0.5 mM EDTA. Peak fractions were concentrated using a 100 kDa MWCO concentrator (Millipore, Cat. No. UFC910008) to ∼5 mg/mL for cryo-EM analysis.

### Cryo-EM sample preparation and data collection

Samples were evaluated by negative-stain EM^37^, prior to cryo-EM grid preparation. Concentrated samples were used for cryo-EM grid preparation. All vitrification steps were carried out on a Vitrobot Mark IV (Thermo Fisher Scientific) at 100% humidity and 8 ℃. A 3.5 µL aliquot of sample was applied onto glow-discharged 300-mesh R1.2/R1.3 UltrAuFoil holey gold grids. After blotting, grids were plunged-frozen in liquid ethane and then transferred to liquid nitrogen for storage. Grids were initially screened using a 200 keV GlaciosCryo-TEM (Thermo Fisher Scientific). Suitable grids showing optimal ice thickness and particle distribution were then transferred to a Titan Krios transmission electron microscope (Thermo Fisher Scientific) operated at 300 keV and equipped with BioQuantum K3 energy filter and a Gatan K3 Direct Electron Detector controlled by EPU software. For all datasets collected in this study, each movie comprised 50 frames, with a total dose of 50 e-/Å2, a calibrated pixel size of 0.92 Å and a defocus range of -0.8 to -1.8 µm.

### Cryo-EM data processing

A total of 6,008 movie micrographs were collected for the GIRK1/2 sample without activator. Each of patch motion correction and patch CTF estimation was performed in cryoSPARC^38^. Dose-weighted micrographs were used for particle picking, followed by multiple rounds of 2D classification to remove contaminants and junk particles. Classes exhibiting clear GIRK channel features were selected for ab initio reconstruction and heterogeneous refinement, which separated particles into a docked-state class and a fully undocked class. The docked-state particles were subjected to nonuniform refinement and local refinement. Aligned particles were then exported from CryoSPARC and imported into RELION^39^ for C4 symmetry expansion, generating all symmetry-related particle orientations. Expanded particles were further classified into eight 3D classes to resolve subtle alignment and conformational differences. Well-aligned classes were retained for a global reconstruction, followed by additional 3D classifications using fixed orientations to isolate finer conformational states. A fully docked class containing 60,413 particles was subjected to CTF refinement, reference-based motion correction, and sharpening to yield a final map at 2.58 Å resolution. A partially docked class containing 36,541 particles underwent an analogous workflow and yielded a final reconstruction at 2.80 Å resolution. The undocked particles obtained from heterogeneous refinement underwent multiple rounds of ab initio reconstruction and classification to remove heterogeneous species. Focused refinement and focused classification on the TMD were then performed to improve alignment in this region. After C4 symmetry expansion on the TMD and further multi-reference 3D classification, the best-aligned class was selected for final TMD reconstruction. Angular information from the TMD refinement was subsequently used to perform local refinement of the CTD, yielding a composite TMD–CTD reconstruction with 3.35 Å (TMD) and 3.62 Å (CTD) resolutions. For the ML297-bound GIRK1/2 dataset, 8,521 movie micrographs were collected and processed using the same procedures for motion correction, CTF estimation, particle picking, and iterative 2D classification. Particles with clear channel features were selected for ab initio reconstruction and heterogeneous classification. The best class containing bound ML297 was refined to a 2.74 Å reconstruction using 79,803 particles. All reported resolutions are based on the gold-standard Fourier shell correlation (FSC) 0.143 criterion as implemented in CryoSPARC. Detailed workflows and intermediate classifications are shown in Figures S2 and S5.

### Model building and refinement

Initial structural models of GIRK1 and GIRK2 predicted by AlphaFold^40^ were obtained from the UniProt database. These models were first rigid-body fitted into the final cryo-EM density map using UCSF Chimera^41^, followed by manual adjustment in Coot^42^ and subsequent real-space refinement in Phenix^43^. The model quality was evaluated using MolProbity^44^, and the refinement statistics are provided in Table S1.

### Pore analysis and visualization

Pore volumes were calculated using MOLEonline^45^ and pore-radius profiles along the permeation pathway were determined using HOLE^46^. All structural visualization and figure rendering were performed in UCSF Chimera and ChimeraX^47^.

### Total internal reflection fluorescence microscopy

GIRK1 and GIRK2 were tagged with mTFP1 (GIRK1-TFP) and TagRFP-T (GIRK2-RFP), respectively, and imaged at the surface of live Xenopus oocytes by TIRF microscopy according to our prior methods^48,49^. Cells were studied after 24-32 hours in low-K⁺ recording solution containing (mM): KCl 2, NaCl 96, MgCl2 1, and HEPES 5, buffered to pH 7.4 with KOH. mTFP1 was excited by a 445-nm laser and GIRK2-RFP by a 561-nm laser (Coherent, Santa Clara, CA). The beams were conditioned for coherence with custom built Keplerian beam expanders upstream of laser cleanup filters (445/10 nm and 561/10 nm, respectively). Laser lines were tuned to provide 10 mW of incident light on a micro mirror positioned below a high numerical aperture apochromat objective (60x, 1.5 NA; Olympus, Waltham, MA) mounted on an RM21 microscope frame equipped with a piezo driven nanopositioning stage (MadCity Labs., Madison, WI). The emission of GIRK1-TFP and GIRK2-RFP were isolated from the excitation beam by an exit micro mirror and a ring-diaphragm positioned below the micro mirror assembly. When the fluors were imaged simultaneously, the emission was split by 510-nm dichroic mirrors placed downstream of parallel 480/40 nm and 620/60 nm bandpass filters. The fluorophores were imaged using a back illuminated sCMOS camera (Teledyne Photometrics, Tucson, AZ) controlled by μ-Manager software^50^. Additional lenses were placed downstream of the camera to adjust the final magnification of the emission image to 100 nm per pixel. All filters and mirrors were from Chroma (Bellow Falls, VT). Lenses, pinholes and diaphragms were from ThorLabs (Newton, NJ). TetraSpeck beads (Thermo, Waltham, MA) were routinely imaged to map the sCMOS chip and to calibrate the evanescent field depth to 100 nm. To assess stoichiometry, fluorophores were photobleached by continual excitation. Data were captured as 16-bit movies of up to 600 frames acquired at 1 Hz. Emission from each fluorophore was captured simultaneously to a mapped sCMOS chip, saved as separate image stacks and processed in an identical manner. Images were corrected for background fluorescence by subtracting the mean of 5 fully bleached frames from the end of each analyzed stack^51^. Fluorescent particles were defined as a discrete 3 x 3-pixel region around a pixel of maximum intensity, as before, and misalignment was corrected in ImageJ using StackReg. Colocalization of partner pixels from the two stacks of images was defined as the presence of both fluorophores with at least 30% of the maximum fluorescence recorded in that region of interest, as before^49^.

### Assessment of surface expression of fluorescent constructs

TIRF microscopy was used to selectively illuminate membrane-proximal fluorescence. HEK293T cells were seeded on 15 mm #1.5 coverslips and transfected 48 hours before imaging. Cells were transfected with either 1 μg GIRK1 WT plus 1 μg GIRK2 WT or mutant, or 1 μg GIRK2 WT plus 1 μg GIRK1 WT or mutant as specified. All GIRK1-TFP constructs and all GIRK2-RFP constructs transfected in HEK293 cells and coverslips were exposed to co-incident laser beams at wavelengths of 445 nm to excite mTFP1 and 561 nm to excite TagRFP-T. Laser lines were tuned to provide 10 mW of incident light on a micromirror positioned below a high numerical aperture apochromat objective (60x, 1.5 numerical aperture; Olympus) mounted on an RM21 microscope frame equipped with a piezo-driven nanopositioning stage (Mad City Labs, Inc). The fluorescence emission of each fluor was isolated from the excitation beam by an exit micromirror and a ring diaphragm positioned below the micromirror assembly. A beam splitter assembly was used to separate the fluorescence emissions of both fluorophores; RFP was imaged through 620/60 nm bandpass filters, and TFP through 480/40 nm bandpass filters, using a back-illuminated sCMOS camera (Teledyne Photometrics) controlled by μManager freeware (University of California San Francisco). The mean fluorescence intensities of selected regions of interest (ROIs) from both bandpass filters were measured to characterize the membrane-proximal expression of both fluorophore-tagged constructs in visibly fluorescent cells in all conditions. ROIs were identified algorithmically using a custom ImageJ Macro script to identify fluorescent regions of cells following background subtraction with a rolling ball radius of 20 pixels. ROIs identified by the algorithm that did not correspond to cell membrane structures upon visual assessment were excluded from analysis.

### Particle classification and scoring

Single-particle TIRF images were analyzed to identify GIRK1-TFP (cyan), GIRK2-RFP (red), and colocalized (overlap of both fluorescence channels) particles in each field of view. Fluorescence-versus-time records for individual particles were scored by manual review as having k₁ distinct mTFP1 bleaching steps and k₂ distinct TagRFP-T bleaching steps. Step counts were assigned based on the number of discrete, monotonic fluorescence transitions visible above background, with a check that successive steps were of similar magnitude. Particles with k₁ ≥ 1 and k₂ ≥ 1 were classified as heteromers. Within the heteromer class, particles displaying exactly one GIRK1-TFP step (k₁ = 1) were analyzed for GIRK2-RFP step counts, as expected for the canonical 1 GIRK1 : 3 GIRK2 stoichiometry.

### Zero-truncated binomial model

Under the standard photobleaching framework^52^, a tetrameric channel containing “n” fluorescently labelled subunits with per-fluorophore step-detection probability p produces a binomial distribution of observed step counts. Because particles can only be detected if at least one step is observed (k ≥ 1), the relevant distribution is the zero-truncated binomial:

P(K = k | K ≥ 1; n, p) = C(n, k) pᵏ (1−p)^(n−k) / [1 − (1−p)ⁿ], k = 1, 2, …, n where C(n, k) = n! / [k!(n−k)!] is the binomial coefficient. The numerator gives the standard binomial probability of observing k steps from n labelled subunits with detection probability p; the denominator removes the probability mass of the unobserved zero-step outcome and renormalizes the distribution. The detection probability p reflects the probability that a single bleaching event is correctly resolved as a discrete step against background noise and combines the underlying fluorophore quantum yield with collection efficiency, integration time, and the criteria used to score steps; p is fit from the data rather than assumed.

### Heteromer fit and goodness-of-fit

For each cRNA mixing ratio, the observed step-count distribution for GIRK2-RFP in the heteromer class (k₁ = 1) was fit to a zero-truncated binomial with n = 3, by maximizing the multinomial log-likelihood with respect to p by bounded numerical optimization in R, following Hines^52^. 95% confidence intervals on p were obtained by parametric bootstrap with 2,000 replicates sampled from the fitted distribution. Goodness-of-fit was assessed by Pearson’s χ² statistic on the three observed bins (k₂ = 1, 2, 3), with one residual degree of freedom (three bins minus one estimated parameter minus one constraint that the bin counts sum to n). The same procedure was applied to the pooled dataset across all seven cRNA conditions. The analysis framework follows Hines^52^ and is consistent with our recent application to Kir2.1–SUMO1 stoichiometry ^53^.

### GIRK2-alone positive control

As a positive control on a known-tetrameric population, GIRK2-TagRFP-T was expressed in *Xenopus* oocytes by injection of GIRK2-RFP cRNA alone, with water substituted for GIRK1 cRNA. The experiment was performed on separate days and using separate biological batches of oocytes from the GIRK1/2 co-injection experiments. GIRK2-RFP-positive membrane particles (k₁ = 0, k₂ ≥ 1) were scored as above and the pooled step distribution was fit to a zero-truncated binomial with n = 4. This experiment validates that GIRK2 traffics alone to the plasma membrane as a tetramer and that the photobleaching framework correctly recovers a 4-subunit binomial distribution on a known-tetrameric population.

### GIRK1-alone trafficking observation

As a control for the trafficking properties of GIRK1, GIRK1-TFP cRNA was injected into Xenopus oocytes alone, with water substituted for GIRK2 cRNA, on a separate day and using a separate biological batch from the co-injection and GIRK2-alone experiments. GIRK1-TFP-positive membrane particles (k₁ ≥ 1, k₂ = 0) were tabulated. No quantitative binomial fit is reported for this dataset given the small particle count; the experiment is reported as a qualitative observation that GIRK1 traffics poorly to the plasma membrane in the absence of an obligate partner subunit.

### Two-electrode voltage clamp

The plasmid DNA for each channel construct was linearized prior to in vitro transcription and transcribed using an mMessage mMachine T7 transcription kit (Thermo Fisher Scientific). *Xenopus* oocytes were surgically extracted in accordance with an IACUC protocol at Northeastern University. Oocytes were then dissociated and defolliculated by collagenase treatment (10 mL of ∼ 1 mg/mL OR2 solution (mM): KCl 2, NaCl 82.5, MgCl_2_ 1, HEPES 5 and pH 7.4 by NaOH) and microinjected with 50 nL of a RNase/DNase free water solution per oocyte containing 1-2 ng each of GIRK1 and GIRK2 cRNAs. Injected oocytes were incubated for 1-2 days in OR2 solution containing 1.8 mM CaCl_2_, 100 units/mL penicillin and 100 µg/mL streptomycin at 17 °C before currents were recorded. Electrodes were pulled using a Flaming-Brown micropipette puller (Sutter Instruments) and were filled with 3 M KCl in 1.5% (w/v) agarose to give resistances between 0.5 and 1.0 MΩ. The oocytes were bathed in ND96 recording solution comprised of (mM): KCl 2, NaCl 96, MgCl_2_ 1, CaCl_2_ 1.8 and HEPES 5, buffered to pH 7.4 with KOH. Where indicated, K^+^ currents were assessed in a high-K^+^ recording solution comprised of (mM): KCl 96, NaCl 2, MgCl_2_ 1, CaCl_2_ 1.8 and HEPES 5, buffered to pH 7.4 with KOH. Currents were measured from whole oocytes using an OC-725D amplifier (Warner Instruments), controlled via a USB interface (National Instruments) and were recorded using WinWCP software (University of Strathclyde). To evoke currents, oocytes were held at 0 mV, stepped to -80 mV for 50 ms, ramped linearly from -80 to +80 mV, held at +80 mV for 50 ms, and finally stepped back to 0 mV. The voltage protocol was repeated every second. At the end of the study, currents were blocked by 10 mM BaCl_2_. The Ba^2+^-sensitive component of the current was analyzed. Typically, a total of 10–15 oocytes from two different *Xenopus* frogs were studied per experiment.

### Gating molecular dynamics simulations

Molecular dynamics simulations were carried out on WT-GIRK1/2, WT-GIRK2, GIRK2-FD mutant, and ML-GIRK1/2 systems in Amber22 using ff19SB forcefield for proteins and LIPID21ff^54^ parameters for lipids. Potassium chloride (150 mM) as well as neutralizing counterions were added to the systems. CHARMM-GUI Membrane Builder web server was used to generate POPC:POPE:POPS:cholesterol (25:5:5:1) lipid bilayer for the systems. TIP3P water box was used to solvate the complexes. The system was built using the tleap module in Amber, followed by hydrogen-mass repartitioning (HMR) to enable a 3-fs integration time step during the production run. Long-range electrostatics were treated with the particle-mesh Ewald (PME) and periodic boundary conditions. Energy minimization was performed by keeping the solute restrained with a harmonic force constant of 500 kcal mol⁻¹ Å⁻² for 100 steps followed by 2,000 steps of steepest descent. The system was heated gradually from 0 K to 100 K over 2,500 steps using Langevin thermostat under constant volume while keeping proteins and lipids weakly restrained (500 kcal mol⁻1 Å⁻2 for the solute; 10 kcal mol⁻1 Å⁻2 for lipids). Subsequently, the temperature was gradually increased from 100 K to 303 K over 50,000 steps using anisotropic pressure coupling to reach 1 atm. The systems were equilibrated at 303 K under NPT conditions using Berendsen barostat and Langevin thermostat. SHAKE constraints were applied to all bonds and a time step of 1 fs was used. This was followed by a series of ten consecutive equilibration runs of 500 ps each (5 ns total) conducted at 303 K using a 2-fs time step. Weak restraints (10–20 kcal·mol⁻1·Å⁻2) were applied to protein backbone to preserve stable membrane-protein packing during equilibration. Each of these equilibrated systems WT-GIRK1/2 and ML-GIRK1/2 were subjected to 10.5 µs and 13 µs of production run, respectively, under NVT conditions. A static electric field with the magnitude of 0.138 kcal·mol⁻1·Å⁻1 (0.06 V/nm, 200 mV across membrane) was applied along the negative z-direction. The results of the simulations were analyzed using various tools, including ChimeraX, VMD^55^ and in-house scripts.

### Interaction pattern analysis

CPPTRAJ and custom Python-based analysis scripts (written in pandas, numpy, and matplotlib) were utilized to explore key ligand-residue interactions throughout the simulation runs. Pairwise hydrogen-bond distances were calculated between critical protein residues and ligand donor/acceptor atoms. These included interactions involving Glu (E141.1), Asp (D173.1), and Trp (W95.1). The hydrogen-bond distances were monitored between ligand amide NH groups and carboxylate oxygens of Asp and Glu and ligand oxygen of carbonyl with Trp. The contact was considered as a hydrogen bond when the donor-acceptor distance was ≤ 3.5 Å. Permeation events were also recorded and mapped with the interaction pattern data to compute total simulation time and frequency of ion permeations associated with each ligand interaction pattern.

### Hydrogen-bond occupancy and ion-permeation counting

For hydrogen bond occupancy (%) analysis, VMD was utilized and total hydrogen bonds of five residue pairs (GIRK2 numbers: Q176.2-E152.2, Q176.2-S148.2, E152.2-T105.2, Y102.2-T105.2, Y102.2-E152.2) were calculated and summed up. The results reported in Figure S10 are averages of 21 individual replicate runs. To calculate the total ion-permeation count, an in-house script was used. K+ ions permeating through the selectivity filter, HBC gate, Upper and Lower G-loop gate were counted in each replicate and averages of 21 replicas are reported. Statistical significance was assessed by t-test. Differences between GIRK2-4KFM and GIRK2-FD were significant (P ≤ 0.01), as were differences between GIRK2-4KFM and GIRK1/2-WT (P ≤ 0.02).

## Data availability

The cryo-EM density maps and corresponding atomic coordinates have been deposited in the Electron Microscopy Data Bank (EMDB) and the PDB, respectively, under the following accession codes: EMD-74962 and 9ZYQ (GIRK1/2 undocked state); EMD-74961 and 9ZYP (GIRK1/2 partially docked state); EMD-74960 and 9ZYO (GIRK1/2 fully docked state); and EMD-74959 and 9ZYN (ML297-bound GIRK1/2 fully docked state). Source data are provided with this paper.

## Acknowledgements

This work was supported by NIH R01 grants HL059949-28 and NS131467-03 to DEL. Thanks to: Heikki Vaananen for EP technical guidance and admin support, Stelios Smirnakis and Mahmoud Nasr at Brigham & Women’s Hospital at Harvard Medical School, Sami Noujaim at the University of South Florida and Satya Kunapuli at Temple University, Logothetis/Plant lab members for discussions, Zhaowen (Norman) Luo for animation preparation. The computations were supported by the ITS (Information Technology Services) Research Computing at Northeastern University and the Argonne Leadership Computing Facility (ALCF) at Argonne National Laboratory.

## Author contributions

L.W., T.K., D.E.L., G.S., L.P., and M.C. designed the study. L.W. performed cryo-EM sample preparation, data collection, processing, model-building and structural analysis. T.K. purified proteins, prepared all mutants, with assistance from N.A. L.P. and J.P. performed single-molecule TIRF experiments and R.C.K. performed the TIRF surface localization experiments. A.S.-C., and E.D. carried out patch-clamp and A.Z., K.W.C. and C.V.-G. TEVC recordings. N.P., M.Z., and Y.L. performed, and M.C. supervised gating molecular dynamics simulations and calculations. S.L. contributed to the structural analysis. D.E.L. and G.S. conceived and supervised the study. L.W. and L.D.P. prepared the figures. L.W., D.E.L., and G.S. wrote the manuscript with input from all authors.

## Extended data figure legends

**Extended Data Figure 1.** Construct design, sequence comparison, purification and representative cryo-EM data for the GIRK1/2 heterotetramer. a, Sequence alignment of human GIRK2 and mouse GIRK2. Non-conserved residues are located at the extreme N- and C-terminal regions (highlighted in yellow); unresolved terminal segments are indicated with thick dashed outlines. b, Schematic of the GIRK2-FLAG-P2A-GIRK1-GFP-His₁₀ construct used for co-expression of GIRK2 and GIRK1 subunits. TEV-cleavable sites were inserted between each gene and its adjacent tag. c, Size-exclusion chromatography of the purified GIRK1/2 complex showing a major tetramer peak (shaded). The corresponding SDS–PAGE gel of the peak fraction confirms the presence of both GIRK1 and GIRK2 subunits. d, Representative cryo-EM micrograph and selected 2D class averages showing high particle quality and clear structural features.

**Extended Data Figure 2.** Cryo-EM processing and reconstruction of GIRK1/2 heterotetramer. a, Cryo-EM data processing workflow for the GIRK1/2 complex. b-d, Final reconstructions of the fully docked (b), partially docked (c) and undocked (d) states. For the undocked state, the TMD and CTD were refined separately and subsequently combined into a composite map. e-h, Gold-standard Fourier shell-correlation (GSFSC) curves and corresponding angular distribution plots for the fully docked state (e), partially docked state (f), and the two components of the undocked state (g and h). Final map resolutions were estimated using the FSC = 0.143 cutoff.

**Extended Data Figure 3.** Agreement between cryo-Cryo-EM density and model. a, Cryo-EM density and model for TM1, TM2, PIP₂ and cholesterol in the 4×PIP₂-bound GIRK1/2 structure. b, Cryo-EM density and model for TM1, TM2, PIP₂ and cholesterol in the 3×PIP₂-bound GIRK1/2 structure. c, Cryo-EM density and model for TM1 and TM2 in the GIRK1/2 undocked state structure. d, Cryo-EM density and model for TM1, TM2, PIP₂ and cholesterol in the ML297-bound GIRK1/2 structure.

**Extended Data Figure 4.** Top-view cryo-EM density showing subunit-specific extracellular interfaces supporting the 1:3 GIRK1/2 stoichiometry. a-c, Cryo-EM maps of the fully docked GIRK1/2 heterotetramer displayed at 3σ (a), 6σ (b), and 9σ (c) contour levels, viewed from the extracellular side. The dashed circles mark a characteristic inter-subunit interface near the selectivity filter. At this position, the three GIRK2 subunits feature V161, which packs against the neighboring GIRK2-Y159 or GIRK1-Y148 depending on its position. In contrast, the corresponding position in GIRK1 is Y150, enabling a larger aromatic side-chain interaction with its adjacent subunit. These subunit-specific features reproduce consistently across all three contour levels, demonstrating the robust alignment and well-resolved density that allow unambiguous identification of GIRK1 relative to the three GIRK2 subunits. d, Overall cryo-EM map and representative local densities for residues that distinguish GIRK1 from GIRK2. GIRK1 is colored pink and GIRK2 subunits are colored in shades of blue/cyan. Local densities are shown for GIRK1 Y150, S65, S256, M98 and F137, and the corresponding GIRK2 residues V161, Y76, Y267, F109 and S148. The map in (d) is contoured at 5σ. These density features support assignment of one GIRK1 subunit and three GIRK2 subunits in the fully docked GIRK1/2 channel.

**Extended Data Figure 5.** Cryo-EM processing and reconstruction of the ML297-bound GIRK1/2 complex. a, Cryo-EM data-processing workflow. b, Final reconstruction of the ML297-bound complex. c, Gold-standard FSC curve; the final resolution was estimated using the FSC = 0.143 criterion. d, Angular distribution plot for the final reconstruction.

**Extended Data Figure 6.** Photobleaching-step distributions support a predominant 1 GIRK1 : 3 GIRK2 heterotetramer across expression ratios. a, GIRK1-mTFP1 and GIRK2-TagRFP-T were co-expressed in *Xenopus* oocytes at the indicated cRNA injection ratios and imaged by TIRF microscopy. For colocalized particles containing one resolved GIRK1-mTFP1 bleaching step, the number of GIRK2-TagRFP-T bleaching steps was quantified. Observed distributions are compared with expected counts from a zero-truncated binomial model assuming three GIRK2 subunits in the heteromer. Across all injection ratios, the data are consistent with a predominant 1 GIRK1 : 3 GIRK2 stoichiometry. b, GIRK2-TagRFP-T expressed alone was used as a positive control for the photobleaching analysis and was fit with a zero-truncated binomial model assuming four GIRK2 subunits. c, GIRK1-mTFP1 expressed alone produced few detectable membrane particles, consistent with poor surface trafficking of GIRK1 in the absence of GIRK2.

**Extended Data Figure 7.** Structural comparison of the GIRK1/2 heterotetramer with the GIRK2 homotetramer. a, Superposition of the fully docked GIRK1/2 heterotetramer (colored by subunit) with the GIRK2 homotetramer (PDB: 6XIT), viewed from the extracellular side. The GIRK1/2 complex exhibits an overall ∼2 Å lateral displacement relative to the GIRK2 homotetramer, reflecting the intrinsic asymmetry of the heterotetrameric assembly. b, Close-up views of the HBC gate (top) and G-loop gate (bottom). In GIRK1/2, the HBC gate shows outward rotation of the gate-forming phenylalanines, and the G-loop gate displays slight opening compared to the corresponding regions in the GIRK2 homotetramer. Arrows indicate the directions of the local conformational differences.

**Extended Data Figure 8.** Structure and sequence comparison between GIRK1 and GIRK2 subunits. a, Structural comparison of GIRK1 (G1s) with each GIRK2 subunit (G2a, G2b, G2c) in the fully docked GIRK1/2 heterotetramer. Superpositions highlight the major divergence between GIRK1 and GIRK2 in the turret, transmembrane helices, and the C-terminal helix, whereas the three GIRK2 subunits are nearly identical to one another. b, Sequence alignment of human GIRK1, human GIRK2, and mouse GIRK2, with secondary structure assignments indicated above the alignment. Regions corresponding to the slide helix, TM1, pore helix, TM2, TMD–CTD linker, and G-loop are annotated. Thick dashed lines mark unresolved regions at the N- and C-termini in the cryo-EM density map.

**Extended Data Figure 9.** Cell surface localization of GIRK1 or GIRK2 mutants co-expressed with the complementary heterotetrameric subunit. HEK293T cells imaged by TIRF microscopy following transfection of the indicated GIRK1-TFP and GIRK2-RFP tagged variants. The mean fluorescence intensities of both fluorophores were measured for all conditions and are presented as mean ± s.e.m. compared to the relative wild-type control from 15–25 cells per condition across 3 biological replicates. For both fluorophores, a consistent level of fluorescence relative to GIRK1 wild type + GIRK2 wild type condition was observed across all constructs. a, Representative fluorescence photomicrographs. b, Summary histogram of GIRK1-TFP fluorescence in the GIRK1/2 channel variants indicated. c, Summary histogram of GIRK2-RFP fluorescence in the GIRK1/2 channel variants indicated.

**Extended Data Figure 10.** Probing the GIRK1 permeation module by mutagenesis of the E141.1 hydrogen-bond network with and without D173.1N. a,b, Sequence alignments of segments of the pore helix containing E141.1 and TM2 containing D173.1 in Kir2 (IRK) and Kir3 (GIRK) channels. c, Representative current traces from wild-type GIRK1/2 (left) and GIRK1(D173N)/2 in low-K⁺ (LK), high-K⁺ (HK) and HK plus Ba²⁺ solutions. d, Basal currents from oocytes expressing G2(S148F) + G2 (1 ng cRNA each) or G2 alone (2 ng cRNA). e, Effects of introducing F137.1S and A94.1T, individually or in combination and with or without D173.1N, on basal G1 + G2 currents.

**Extended Data Figure 11.** GMD links reduced hydrogen-bond occupancy and acidic residues beneath the selectivity filter to increased permeation. a-c, Average hydrogen-bond occupancy (%) in GIRK1/2-WT (a), GIRK2-4KFM (b) and GIRK2-FD (c). d, Total hydrogen-bond occupancy in the homotetrameric and heterotetrameric systems, averaged across 21 replicate runs. e, K⁺ permeation counts for GIRK1/2-WT, GIRK2-4KFM and GIRK2-FD, averaged across 21 replicate runs. P values for the indicated comparisons are shown.

**Extended Data Figure 12.** Interaction patterns and ion-permeation dynamics of ML297-bound GIRK1/2. a, Summary of simulation time (ST) and permeation counts (PC) associated with interactions of ML297 with E141.1, D173.1, W95.1, S177.2 and S181.2. Multiple 500-ns runs yielded >10 μs total simulation time for each system. b-d, Representative ML297 conformations highlighting hydrogen-bond geometry and the dominant interaction patterns sampled in the GIRK1/2 binding pocket.

