## Extended Data for "Asymmetric assembly underlies activation mechanism of neuronal GIRK1/2 channels"

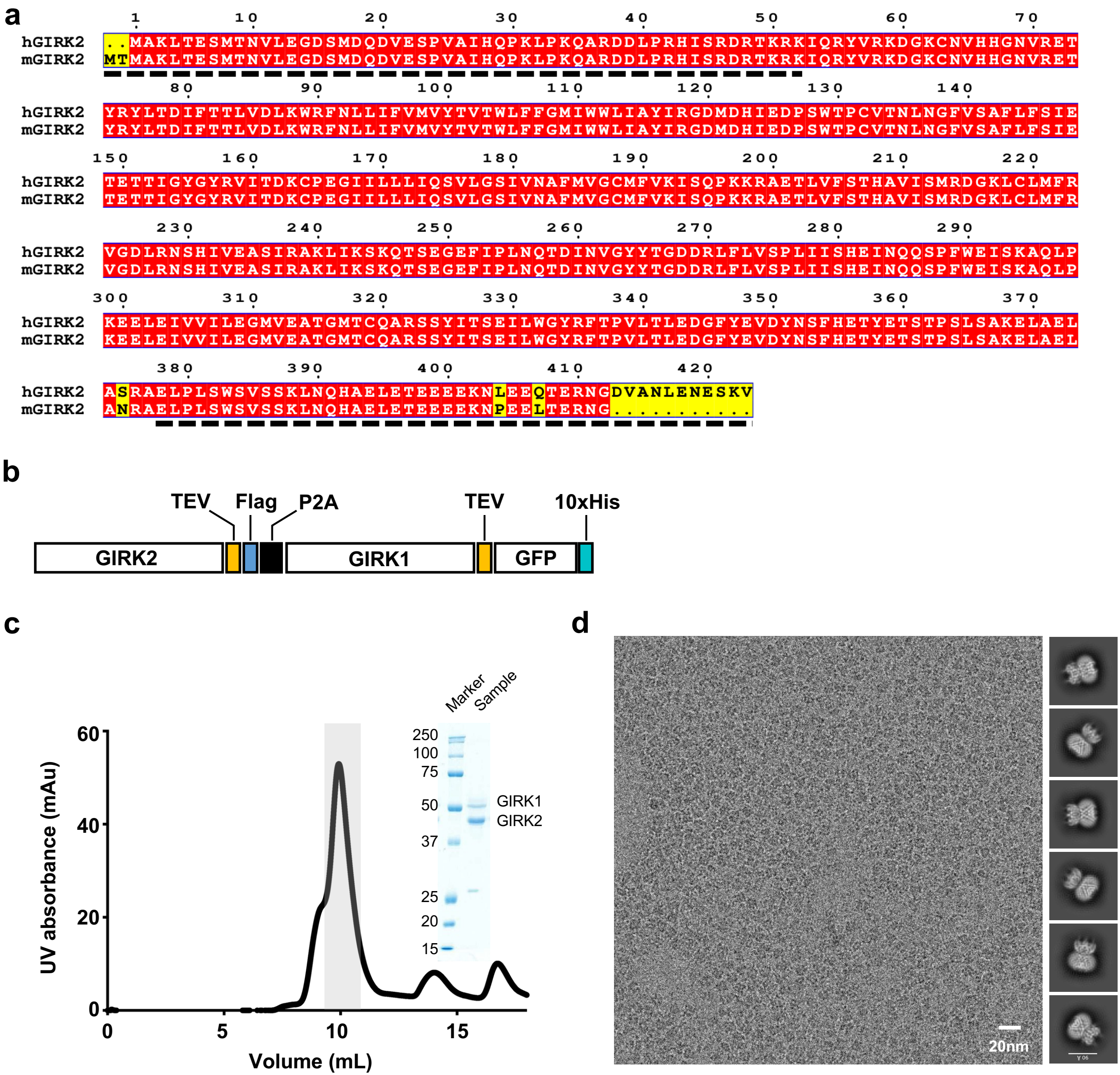
a
1 10 20 30 40 50 60 70
hGIRK2 ..MAKLTESMTNVLEGDSMDQDVESPVAIHQPKLPKQARDDLPRHISRDRTKRKIQRYVRKDGKCNVHHGNVRET
mGIRK2 MTMAKLTESMTNVLEGDSMDQDVESPVAIHQPKLPKQARDDLPRHISRDRTKRKIQRYVRKDGKCNVHHGNVRET
80 90 100 110 120 130 140
hGIRK2 YRYLTDIFTTLVDLKWRFNLLIFVMVYTVTWLFFGMIWWLIAYIRGDMDHIEDPSWTPCVTNLNGFVSAFLFSIE
mGIRK2 YRYLTDIFTTLVDLKWRFNLLIFVMVYTVTWLFFGMIWWLIAYIRGDMDHIEDPSWTPCVTNLNGFVSAFLFSIE
150 160 170 180 190 200 210 220
hGIRK2 TETTIGYGYRVITDKCPEGIILLLIQSVLGSIVNAFMVGCMFVKISQPKKRAETLVFSTHAVISMRDGKLCLMFR
mGIRK2 TETTIGYGYRVITDKCPEGIILLLIQSVLGSIVNAFMVGCMFVKISQPKKRAETLVFSTHAVISMRDGKLCLMFR
230 240 250 260 270 280 290
hGIRK2 VGDLRNSHIVEASIRAKLIKSKQTSEGEFIPLNQTDINVGYYTGDDRLFLVSPLIISHEINQQSPFWEISKAQLP
mGIRK2 VGDLRNSHIVEASIRAKLIKSKQTSEGEFIPLNQTDINVGYYTGDDRLFLVSPLIISHEINQQSPFWEISKAQLP
300 310 320 330 340 350 360 370
hGIRK2 KEELEIVVILEGMVEATGMTCQARSSYITSEILWGYRFTPVLTLEDGFYEVDYNSFHETYETSTPSLSAKELAEL
mGIRK2 KEELEIVVILEGMVEATGMTCQARSSYITSEILWGYRFTPVLTLEDGFYEVDYNSFHETYETSTPSLSAKELAEL
380 390 400 410 420
hGIRK2 ASRAELPLSWSVSSKLNQHAELETEEEEKNLEEQTERNGDVANLENESKV
mGIRK2 ANRAELPLSWSVSSKLNQHAELETEEEEKNPEELTERNG..........
b
TEV
Flag
P2A
TEV
10xHis
GIRK2
GIRK1
GFP
c
Marker
Sample
250
100
75
50
37
25
20
15
GIRK1
GIRK2
UV absorbance (mAu)
60
40
20
0
0
5
10
15
Volume (mL)
d
20nm

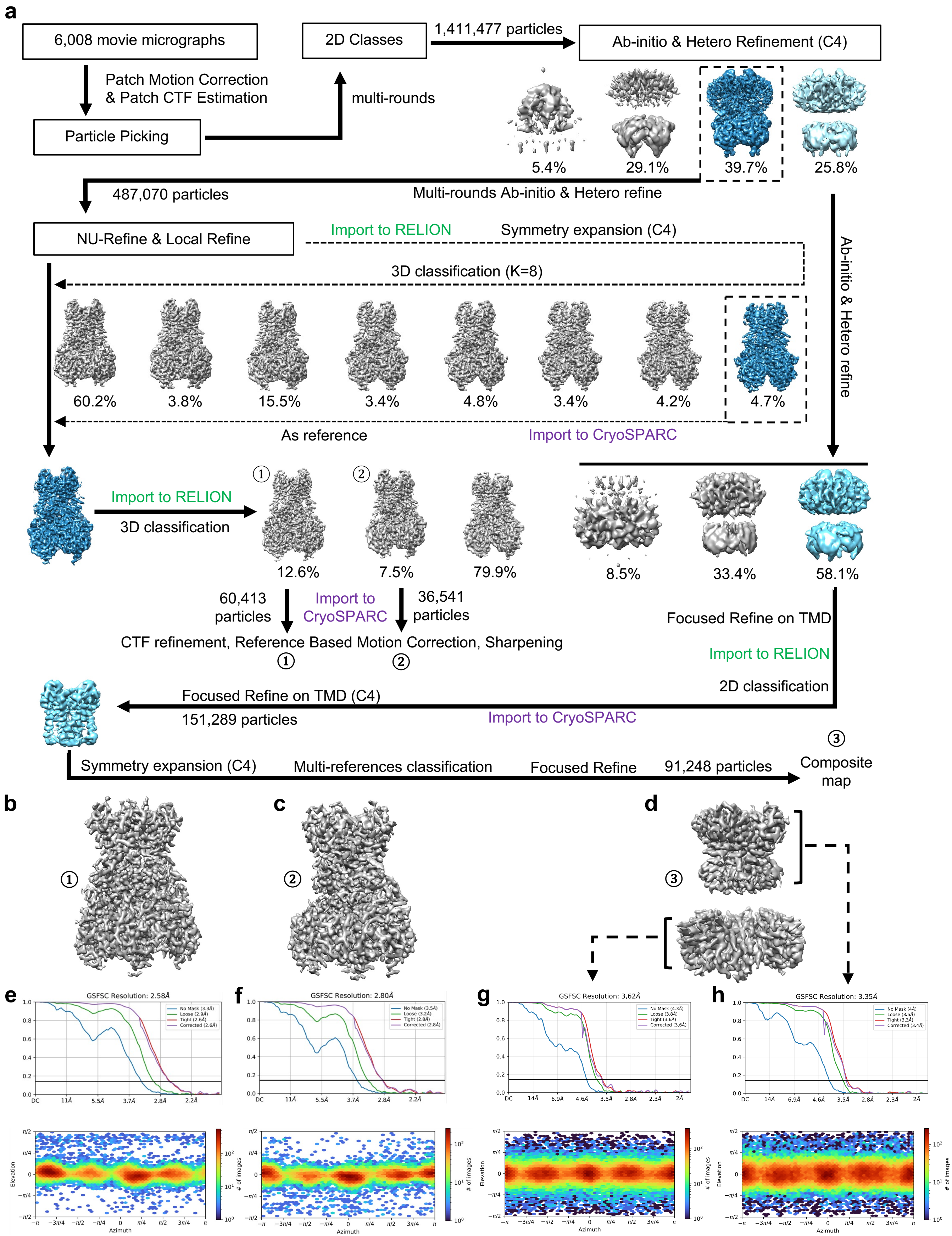
a
6,008 movie micrographs
Patch Motion Correction
& Patch CTF Estimation
Particle Picking
multi-rounds
2D Classes
1,411,477 particles
Ab-initio & Hetero Refinement (C4)
5.4%
29.1%
39.7%
25.8%
487,070 particles
Multi-rounds Ab-initio & Hetero refine
NU-Refine & Local Refine
Import to RELION
Symmetry expansion (C4)
3D classification (K=8)
Ab-initio & Hetero refine
60.2%
3.8%
15.5%
3.4%
4.8%
3.4%
4.2%
4.7%
As reference
Import to CryoSPARC
Import to RELION
3D classification
①
②
12.6%
7.5%
79.9%
8.5%
33.4%
58.1%
60,413 particles
Import to CryoSPARC
36,541 particles
CTF refinement, Reference Based Motion Correction, Sharpening
①
②
Focused Refine on TMD
Import to RELION
2D classification
Focused Refine on TMD (C4)
151,289 particles
Import to CryoSPARC
Symmetry expansion (C4)
Multi-references classification
Focused Refine
91,248 particles
③
Composite map
b
①
c
②
d
③
e
GSFSC Resolution: 2.58Å
f
GSFSC Resolution: 2.80Å
g
GSFSC Resolution: 3.62Å
h
GSFSC Resolution: 3.35Å
Elevation
Azimuth
### of images

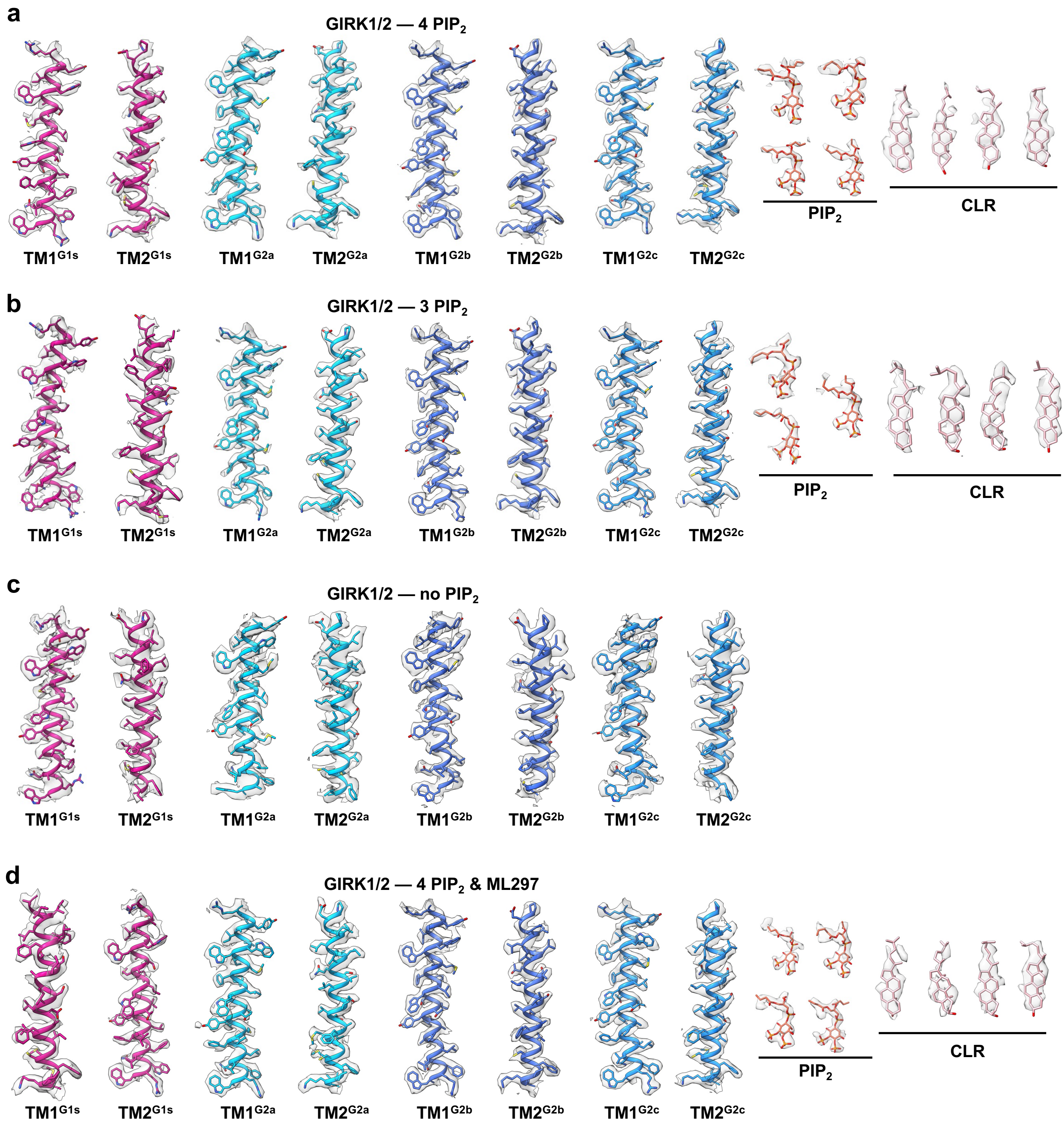
a
GIRK1/2 — 4 PIP2
PIP2
CLR
TM1G1s
TM2G1s
TM1G2a
TM2G2a
TM1G2b
TM2G2b
TM1G2c
TM2G2c
b
GIRK1/2 — 3 PIP2
PIP2
CLR
TM1G1s
TM2G1s
TM1G2a
TM2G2a
TM1G2b
TM2G2b
TM1G2c
TM2G2c
c
GIRK1/2 — no PIP2
TM1G1s
TM2G1s
TM1G2a
TM2G2a
TM1G2b
TM2G2b
TM1G2c
TM2G2c
d
GIRK1/2 — 4 PIP2 & ML297
PIP2
CLR
TM1G1s
TM2G1s
TM1G2a
TM2G2a
TM1G2b
TM2G2b
TM1G2c
TM2G2c

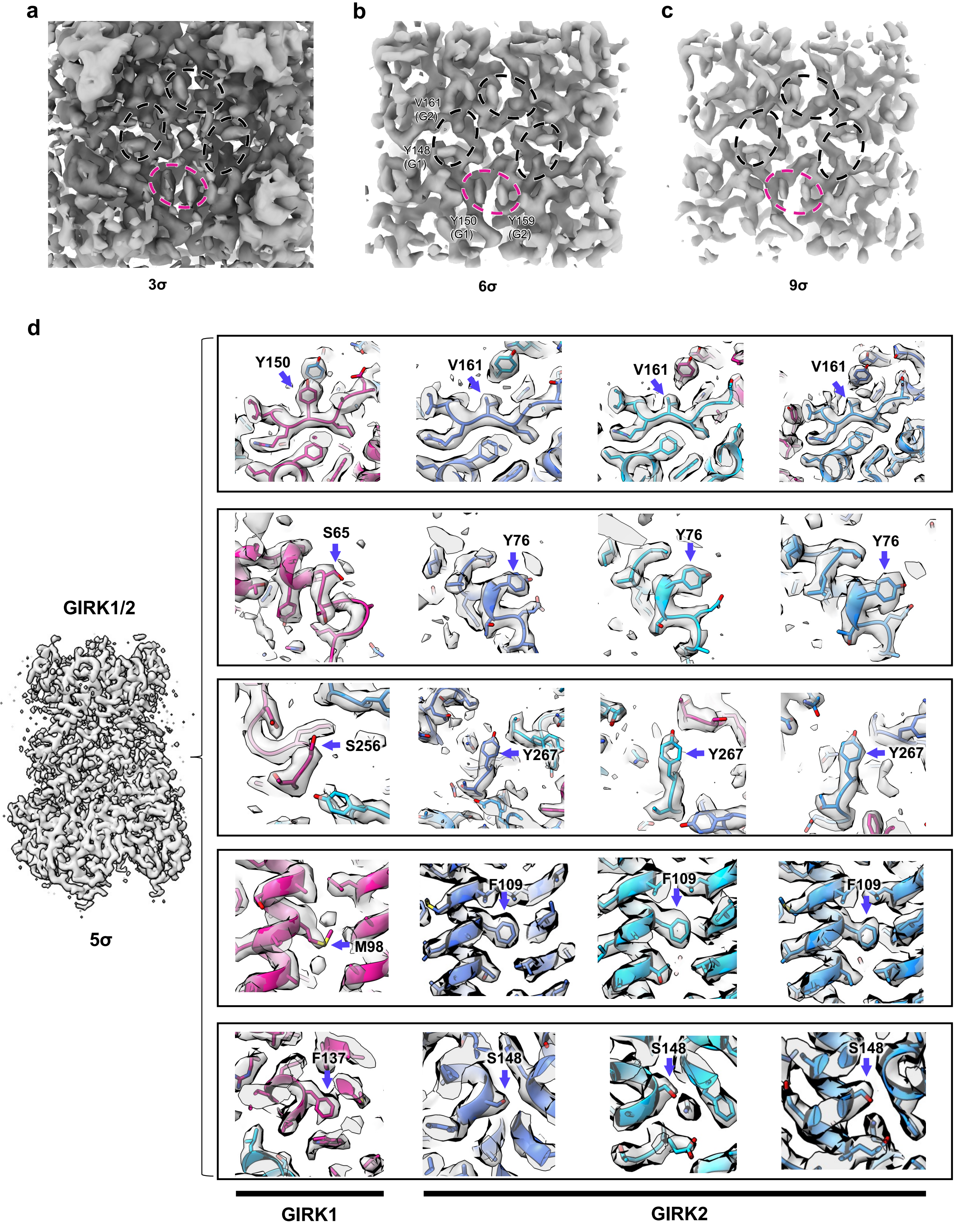

a
b
c
V161
(G2)
Y148
(G1)
Y150
(G1)
Y159
(G2)
3σ
6σ
9σ
d
GIRK1/2
5σ
Y150
V161
V161
V161
S65
Y76
Y76
Y76
S256
Y267
Y267
Y267
M98
F109
F109
F109
F137
S148
S148
S148
GIRK1
GIRK2

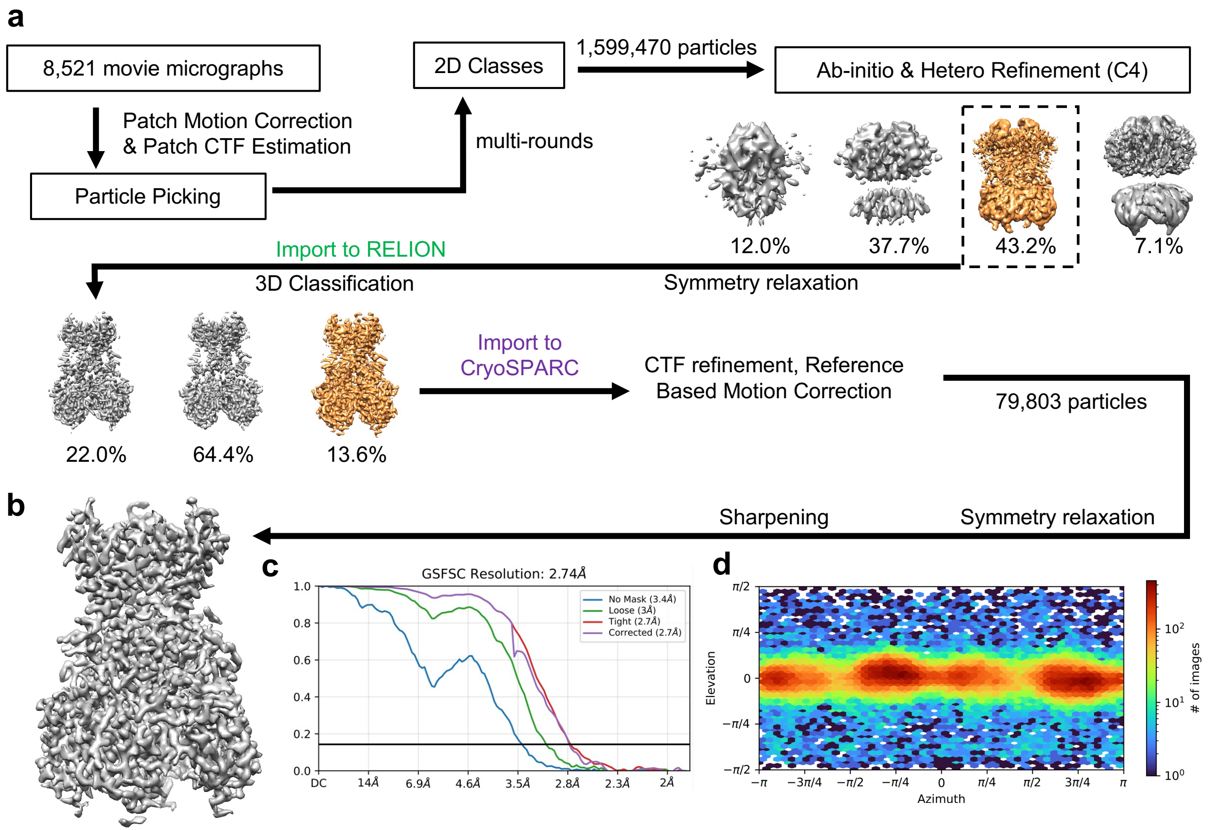
a
8,521 movie micrographs
Patch Motion Correction
& Patch CTF Estimation
Particle Picking
multi-rounds
2D Classes
1,599,470 particles
Ab-initio & Hetero Refinement (C4)
12.0%
37.7%
43.2%
7.1%
Import to RELION
3D Classification
Symmetry relaxation
22.0%
64.4%
13.6%
Import to
CryoSPARC
CTF refinement, Reference
Based Motion Correction
79,803 particles
Sharpening
Symmetry relaxation
b
c
GSFSC Resolution: 2.74Å
No Mask (3.4Å)
Loose (3Å)
Tight (2.7Å)
Corrected (2.7Å)
DC
14Å
6.9Å
4.6Å
3.5Å
2.8Å
2.3Å
2Å
d
Elevation
Azimuth
### of images

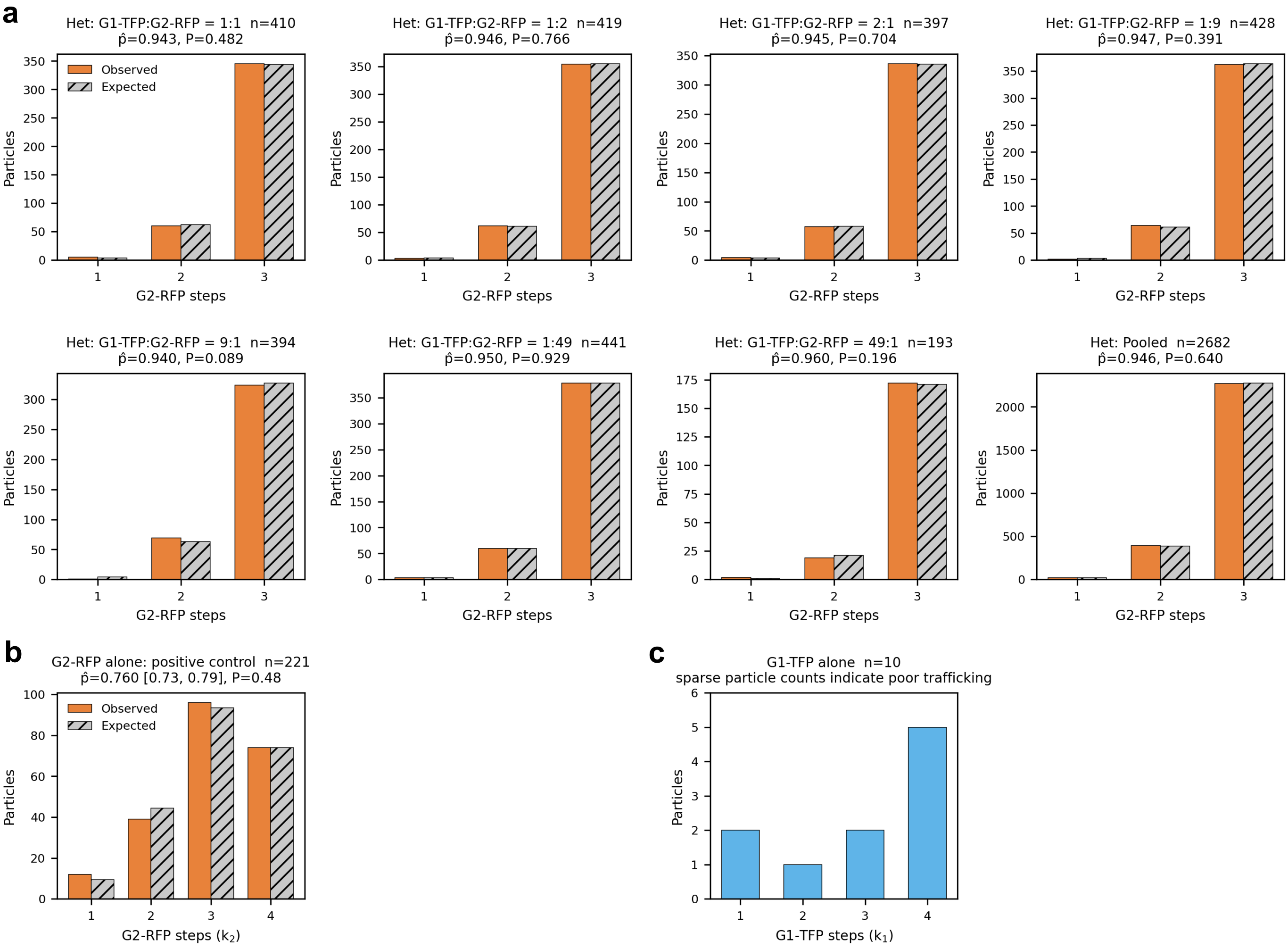
a
Het: G1-TFP:G2-RFP = 1:1 n=410
p̂=0.943, P=0.482
Observed
Expected
Particles
G2-RFP steps
Het: G1-TFP:G2-RFP = 1:2 n=419
p̂=0.946, P=0.766
Het: G1-TFP:G2-RFP = 2:1 n=397
p̂=0.945, P=0.704
Het: G1-TFP:G2-RFP = 1:9 n=428
p̂=0.947, P=0.391
Het: G1-TFP:G2-RFP = 9:1 n=394
p̂=0.940, P=0.089
Het: G1-TFP:G2-RFP = 1:49 n=441
p̂=0.950, P=0.929
Het: G1-TFP:G2-RFP = 49:1 n=193
p̂=0.960, P=0.196
Het: Pooled n=2682
p̂=0.946, P=0.640
b
G2-RFP alone: positive control n=221
p̂=0.760 [0.73, 0.79], P=0.48
Observed
Expected
Particles
G2-RFP steps ($k_2$)
c
G1-TFP alone n=10
sparse particle counts indicate poor trafficking
Particles
G1-TFP steps ($k_1$)

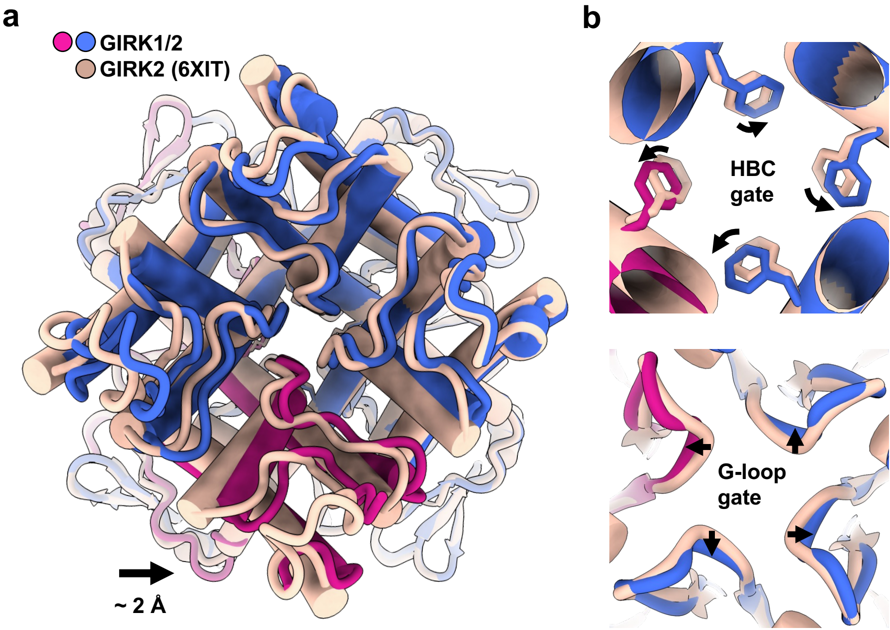
a
GIRK1/2
GIRK2 (6XIT)
~ 2 Å
b
HBC
gate
G-loop
gate

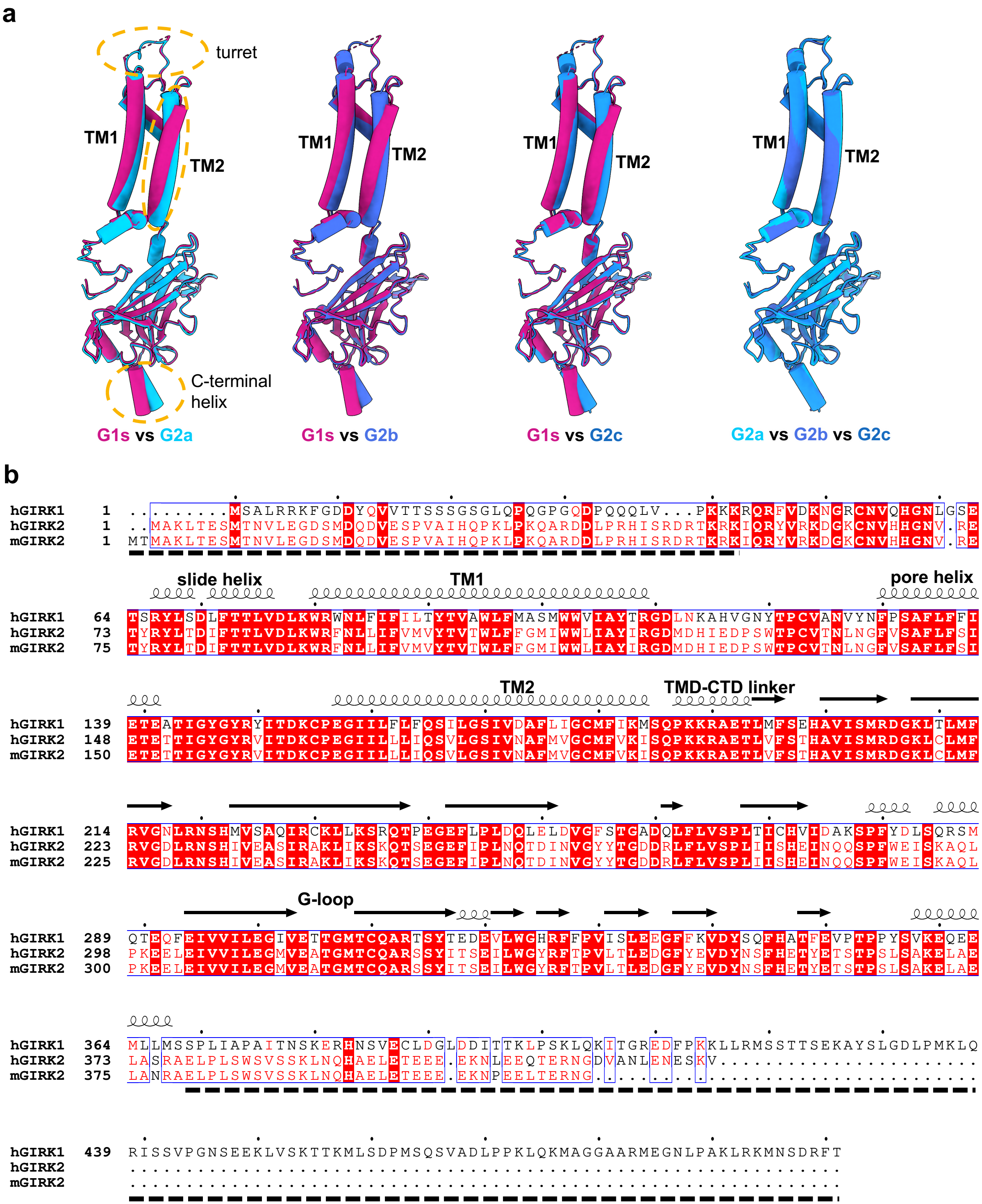
a
turret
TM1
TM2
C-terminal helix
G1s vs G2a
G1s vs G2b
G1s vs G2c
G2a vs G2b vs G2c
b
hGIRK1
hGIRK2
mGIRK2
slide helix
TM1
pore helix
TM2
TMD-CTD linker
G-loop

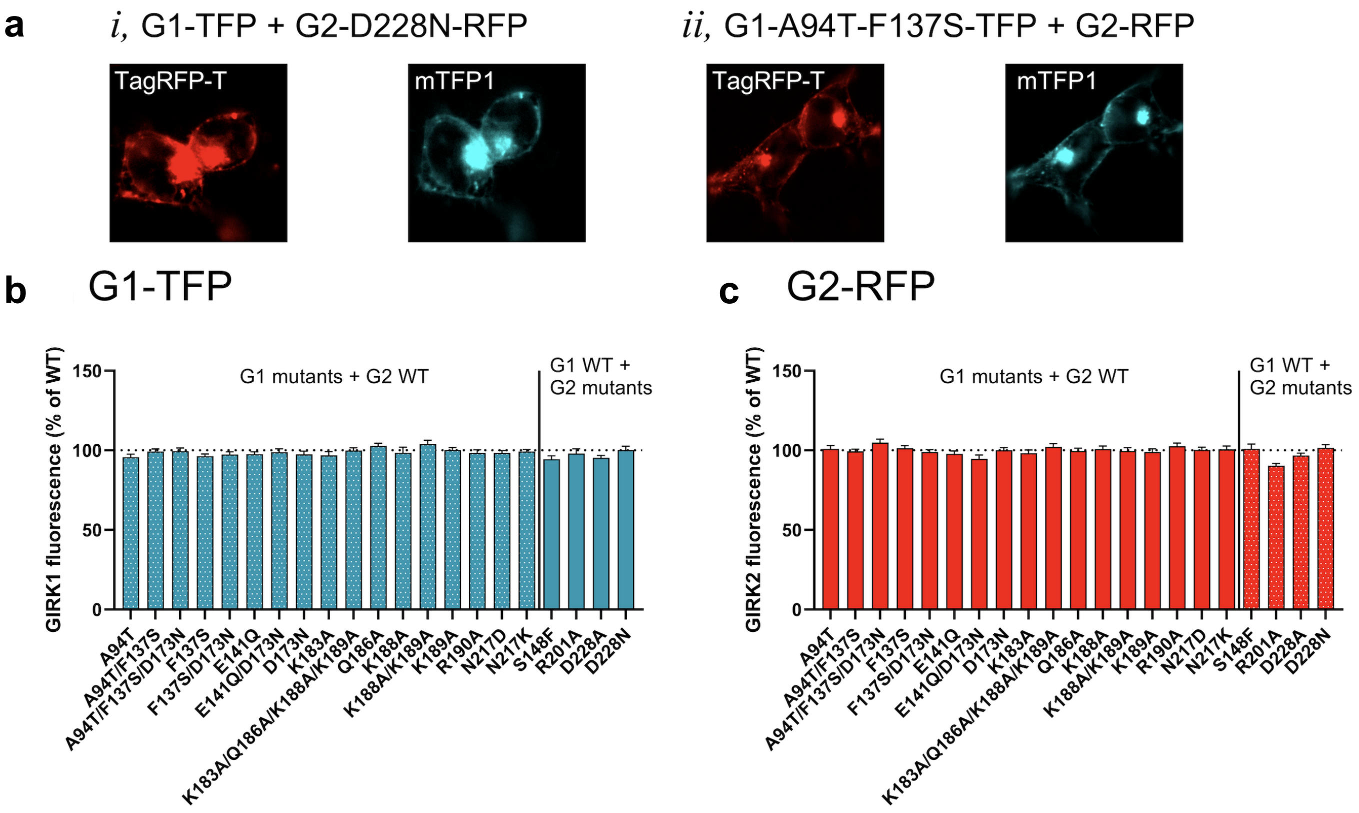
a
i, G1-TFP + G2-D228N-RFP
TagRFP-T
mTFP1
ii, G1-A94T-F137S-TFP + G2-RFP
TagRFP-T
mTFP1
b
G1-TFP
GIRK1 fluorescence (% of WT)
G1 mutants + G2 WT
G1 WT + G2 mutants
0
50
100
150
A94T
A94T/F137S
A94T/F137S/D173N
F137S
F137S/D173N
E141Q
E141Q/D173N
D173N
K183A
K183A/Q186A/K188A/K189A
Q186A
K188A
K188A/K189A
K189A
R190A
N217D
N217K
S148F
R201A
D228A
D228N
c
G2-RFP
GIRK2 fluorescence (% of WT)
G1 mutants + G2 WT
G1 WT + G2 mutants
0
50
100
150
A94T
A94T/F137S
A94T/F137S/D173N
F137S
F137S/D173N
E141Q
E141Q/D173N
D173N
K183A
K183A/Q186A/K188A/K189A
Q186A
K188A
K188A/K189A
K189A
R190A
N217D
N217K
S148F
R201A
D228A
D228N

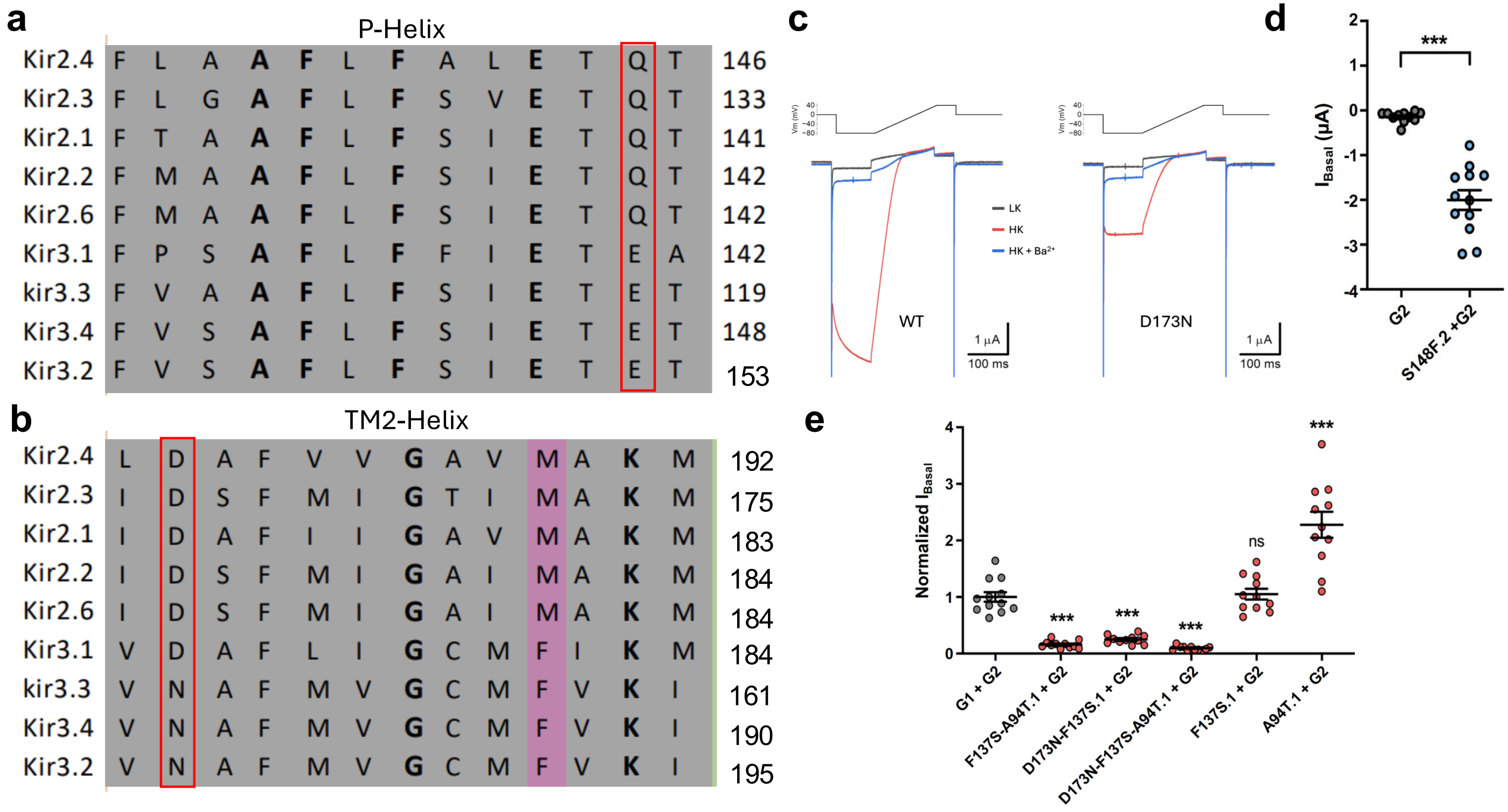

a
P-Helix
Kir2.4 F L A A F L F A L E T Q T 146
Kir2.3 F L G A F L F S V E T Q T 133
Kir2.1 F T A A F L F S I E T Q T 141
Kir2.2 F M A A F L F S I E T Q T 142
Kir2.6 F M A A F L F S I E T Q T 142
Kir3.1 F P S A F L F F I E T E A 142
kir3.3 F V A A F L F S I E T E T 119
Kir3.4 F V S A F L F S I E T E T 148
Kir3.2 F V S A F L F S I E T E T 153
b
TM2-Helix
Kir2.4 L D A F V V G A V M A K M 192
Kir2.3 I D S F M I G T I M A K M 175
Kir2.1 I D A F I I G A V M A K M 183
Kir2.2 I D S F M I G A I M A K M 184
Kir2.6 I D S F M I G A I M A K M 184
Kir3.1 V D A F L I G C M F I K M 184
kir3.3 V N A F M V G C M F V K I 161
Kir3.4 V N A F M V G C M F V K I 190
Kir3.2 V N A F M V G C M F V K I 195
c
LK
HK
HK + Ba2+
WT
D173N
1 μA
100 ms
d
I Basal (μA)
***
G2
S148F.2 +G2
e
Normalized I Basal
***
ns
G1 + G2
F137S-A94T.1 + G2
D173N-F137S.1 + G2
D173N-F137S-A94T.1 + G2
F137S.1 + G2
A94T.1 + G2

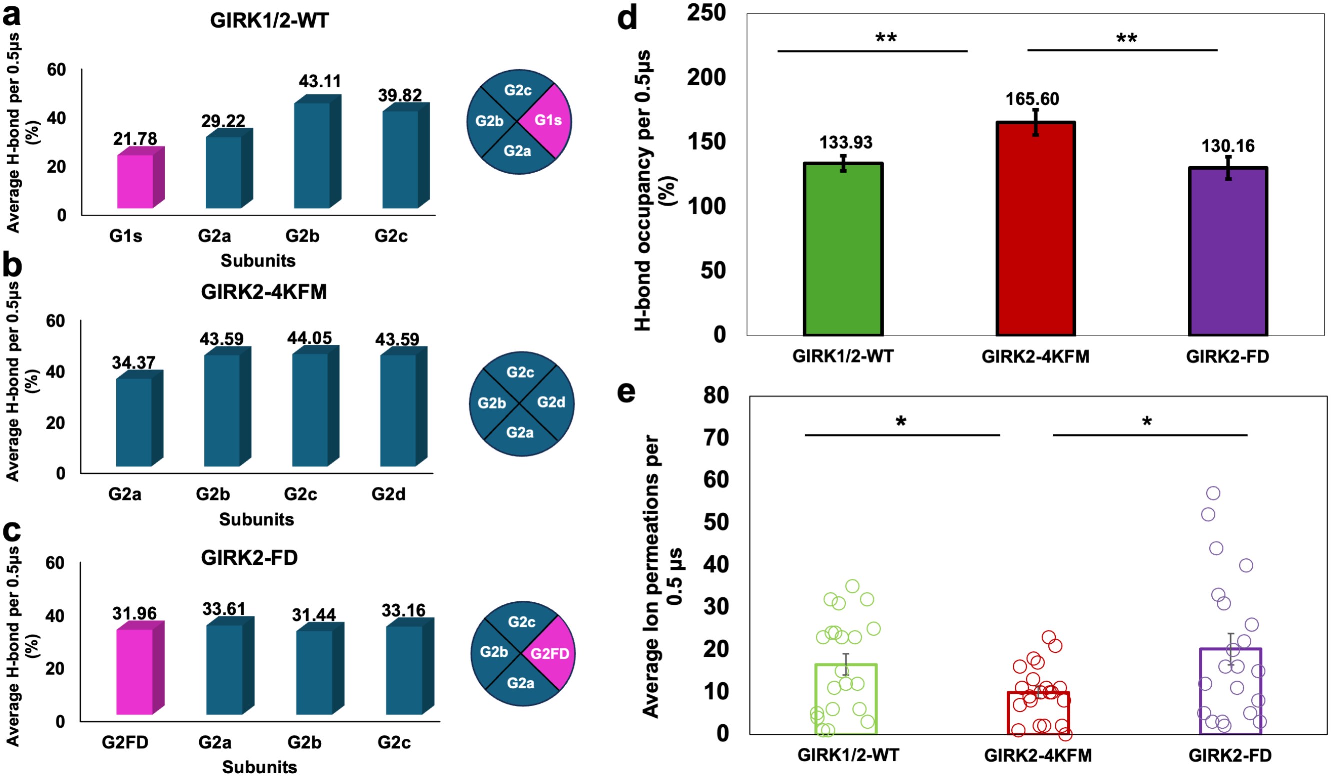

a
GIRK1/2-WT
Average H-bond per 0.5μs (%)
60
40
20
0
21.78
29.22
43.11
39.82
G1s
G2a
G2b
G2c
Subunits
G2c
G2b
G1s
G2a
b
GIRK2-4KFM
Average H-bond per 0.5μs (%)
34.37
43.59
44.05
43.59
G2a
G2b
G2c
G2d
Subunits
G2c
G2b
G2d
G2a
c
GIRK2-FD
Average H-bond per 0.5μs (%)
31.96
33.61
31.44
33.16
G2FD
G2a
G2b
G2c
Subunits
G2c
G2b
G2FD
G2a
d
H-bond occupancy per 0.5μs (%)
250
200
150
100
50
0
**
**
133.93
165.60
130.16
GIRK1/2-WT
GIRK2-4KFM
GIRK2-FD
e
Average Ion permeations per 0.5 μs
80
70
60
50
40
30
20
10
0
*
*
GIRK1/2-WT
GIRK2-4KFM
GIRK2-FD

**a**

| SYSTEMS | Interactions involving residue E141.1/D173.1/W95.1/S177.2/S181.2 | | Total ST (μs) | Total PC (per 0.5 μs) |
|---|---|---|---|---|
| | ST in *ns* | PC | | |
| ML297-G1/2 | E: 7098 (55%)<br>D: 5263 (41%)<br>W: 5321 (41.1%)<br>S177: 1932 (14.9%)<br>S181: 3709 (28.5%) | E: 323 (60%)<br>D: 202 (37%)<br>W: 279 (51.5%)<br>S177: 178 (32.8%)<br>S181: 119 (22%) | 13 | 19 |
| WT-G1/2 | - | - | 10.5 | 17 |

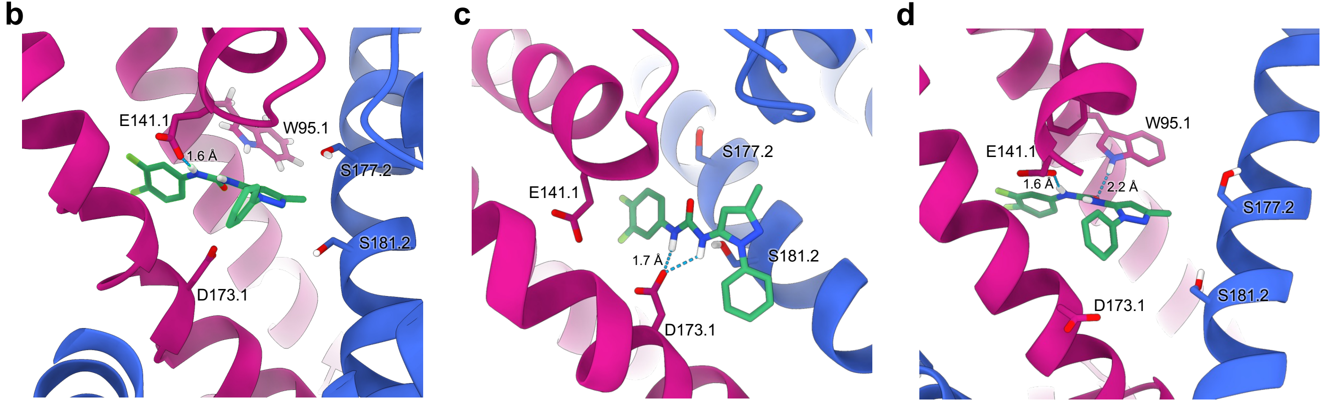


**RUN 17, E (20% ST, 23.8% PC)**

**RUN 7, D (22.1% ST, 10.3% PC)**

**RUN 20, E-W (14.7% ST, 37.5% PC)**

**Extended Data Table 1 Analysis of GIRK subunit stoichiometries**

#### Part A. Heteromer binomial fits (co-injection experiments)

| Condition (cRNA G1:G2) | n | Observed counts ($k_2$=1, 2, 3) | p̂ (95% CI) | Expected counts ($k_2$=1, 2, 3) | $\chi^2$ | P (goodness-of-fit) |
|---|---|---|---|---|---|---|
| 1:1 | 410 | (5, 60, 345) | 0.943 (0.931–0.955) | (3.8, 62.4, 343.8) | 0.493 | 0.483 |
| 1:2 | 419 | (3, 62, 354) | 0.946 (0.934–0.958) | (3.5, 61.0, 354.5) | 0.088 | 0.767 |
| 2:1 | 397 | (4, 57, 336) | 0.945 (0.933–0.957) | (3.4, 58.0, 335.6) | 0.145 | 0.704 |
| 1:9 | 428 | (2, 64, 362) | 0.947 (0.935–0.959) | (3.5, 61.7, 362.7) | 0.735 | 0.391 |
| 9:1 | 394 | (1, 69, 324) | 0.940 (0.926–0.953) | (3.0, 64.1, 326.8) | 2.898 | 0.089 |
| 1:49 | 441 | (3, 60, 378) | 0.950 (0.939–0.961) | (3.0, 59.5, 378.5) | 0.008 | 0.929 |
| 49:1 | 193 | (2, 19, 172) | 0.960 (0.943–0.976) | (1.0, 22.3, 169.8) | 1.669 | 0.196 |
| Pooled | 2682 | (20, 391, 2271) | 0.946 (0.941–0.951) | (22.0, 387.1, 2273.0) | 0.219 | 0.640 |

#### Part B. GIRK2-alone positive control

| Experiment | n | Observed ($k_2$ = 1, 2, 3, 4) | p̂ (95% CI) | Expected ($k_2$ = 1, 2, 3, 4) | $\chi^2$ | P |
|---|---|---|---|---|---|---|
| GIRK2 alone | 221 | (12, 39, 96, 74) | 0.760 (0.732–0.788) | (9.3, 44.3, 93.4, 73.9) | 1.47 | 0.48 |

#### Part C. GIRK1-alone trafficking observation

| Experiment | n | (1,0) | (2,0) | (3,0) | (4,0) |
|---|---|---|---|---|---|
| GIRK1 alone | 10 | 2 | 1 | 2 | 5 |
